# LAMP2A regulates the loading of proteins into exosomes

**DOI:** 10.1101/2021.07.26.453637

**Authors:** João Vasco Ferreira, Ana da Rosa Soares, José Ramalho, Catarina Máximo Carvalho, Maria Helena Cardoso, Petra Pintado, Ana Sofia Carvalho, Hans Christian Beck, Rune Matthiesen, Mónica Zuzarte, Henrique Girão, Guillaume van Niel, Paulo Pereira

**Author notes:** The authors contributed equally to this work (João Vasco Ferreira, Ana da Rosa Soares).

## Abstract

Exosomes are extracellular vesicles of endosomal origin released by virtually all cell types across metazoans. Exosomes are active vehicles of intercellular communication and can transfer lipids, RNAs and proteins between different cells, tissues or organs. However, the mechanisms that regulate the selective loading of cytosolic proteins into these vesicles are still largely unknow. Here we describe a mechanism whereby proteins containing a pentapeptide sequence, biochemically related to the KFERQ-motif, are loaded into a subpopulation of exosomes in a process that is dependent on the membrane protein LAMP2A. Moreover, this mechanism is independent of the ESCRT machinery components TSG101 and VPS4b and dependent on HSC70, CD63, Alix, Syntenin-1, Rab31 and ceramides. The transcription factor and master regulator of hypoxia HIF1A is loaded into exosomes by this mechanism to transport hypoxia signaling to normoxic cells. Additionally, by tagging fluorescent proteins with KFERQ-like sequences we were able to follow inter-organ transfer of exosomes in zebrafish larvae. Our findings identify LAMP2A as a key component in exosome biogenesis while opening new avenues for exosome engineering by allowing the loading of bioactive proteins by tagging them with KFERQ-like motifs.

## Introduction

Exosomes are nanosized vesicles of 40–160 nm in diameter secreted by most cell types to the extracellular space^1^. Exosomes, containing lipids, metabolites, proteins and RNA, can travel and transfer cellular information from one cell to another^1–3^ in diverse biological processes such as immune function, viral infection, metabolic regulation, tumor metastasis and neurodegeneration^4^. Additionally, circulating exosomes have several potential uses, such as disease biomarkers in liquid biopsies, while efforts are being made to create engineered exosomes for a variety of therapeutic approaches, such as vaccines and gene therapy, or to efficiently target undruggable proteins.

Exosomes are formed by the in-budding of the endosomal membrane to create vesicle-laden endosomes, filled with intraluminal vesicles (ILVs), referred to as multivesicular bodies (MVBs)^5^. Fusion of some of these MVBs with the plasma membrane leads to the release of ILVs to the extracellular space as exosomes^5^. Outside the originating cell, exosomes can dock and fuse with (or get internalized by) other cells, to deliver their cargo^5, 6^.

ILV biogenesis was considered to be largely mediated by the ESCRT (endosomal sorting complex required for transport) machinery^1, 5, 7^. Nonetheless, ESCRT depleted cells can still generate ILVs^8^. Alternative mechanisms have been reported to assist in ILV formation and cargo sorting such as the sphingolipid ceramide ^8^, the tetraspanin CD63, the Toll-like receptor trafficking chaperone UNC93B1 or the Syndecan-Syntenin-Alix pathway^1, 9–12^. However, the selection and sorting of soluble cytoplasmic components into exosomes remains largely unknown. Significantly, there is ample evidence suggesting that the cargo repertoire of exosomes does not necessarily reflect the cytosolic contents of the originating cell^13, 14^, indicating the existence of some type of active soluble cargo selection or triage.

In this study we show that proteins containing amino acid sequences biochemically related to the KFERQ-motif^15^, are loaded into a subpopulation of exosomes. We identified the following as essential components of the machinery: the Lysosome-Associated Membrane Protein 2, isoform A (LAMP2A), previously recognized as a protein that targets KFERQ-containing substrates to the lysosome^15^ and the molecular chaperone HSC70 (heat shock 70 kDa protein 8). Additionally, the loading of proteins takes place at the early endosomal membrane and relies on the endosomal proteins CD63, Alix, Syntenin-1 and Rab31, rather than on components of the ESCRT-machinery proteins such as TSG101 and VPS4b. We further show that this mechanism is of biological relevance and occurs in animals. For example, we showed that the hypoxia master regulator HIF1A (hypoxia-inducible transcription factor 1 alpha) is loaded into exosomes by the action of its KFERQ-like motif and the presence of LAMP2A in endosomes, to transfer HIF1A transcriptional activity from hypoxic to normoxic cells, *in vitro* as well as in the zebrafish. Moreover, our findings enabled us to develop tools to track the transfer of exosomes from one tissue to another in zebrafish larvae, creating new possibilities for the study of exosome-mediated inter-organ communication.

## Results

### LAMP2A is required for the sorting of selected proteins into small extracellular vesicles (sEVs)

To investigate the impact of LAMP2A in exosome protein cargo we knocked-out (KO) the LAMP2A gene using CRISPR-Cas9. To expedite the creation of the KO cells we took advantage of a widely used human cell line with normal chromosome number from the retina pigment epithelium (ARPE-19). LAMP2A originates from the alternative splicing of the LAMP2 gene. The LAMP2 gene contains 9 exons, including 3 different splice variants of the exon 9 (A, B and C) (Fig.1A). All splice variants share a common luminal domain but contain different cytosolic and transmembrane regions. We were able to isolate several clones of ARPE-19 cells without LAMP2A while maintaining the expression of other LAMP2 isoforms (SupFig.1A).

**Figure 1.**
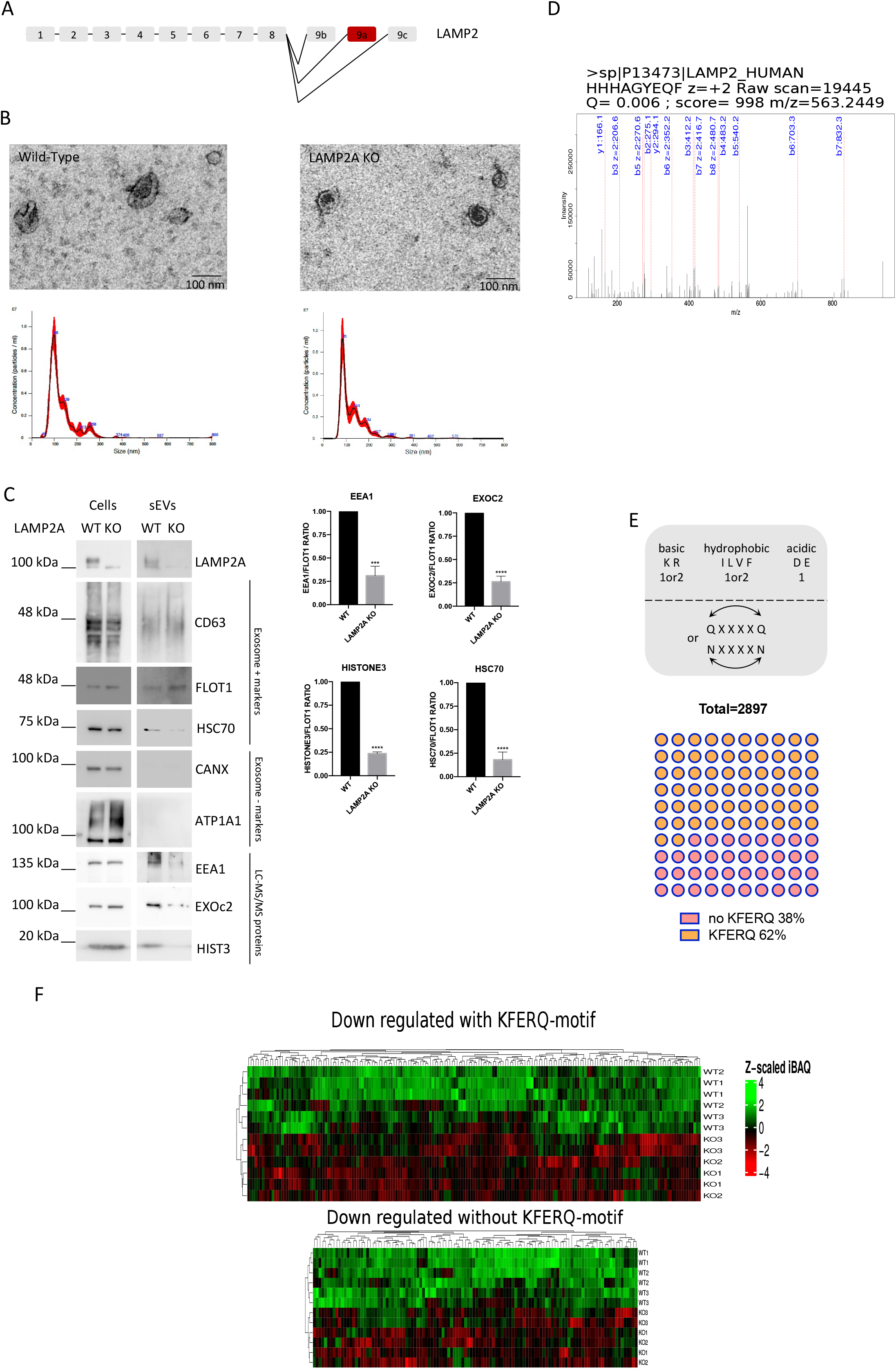
LAMP2A knock-out leads to the downregulation of KFERQ-containing proteins in sEVs. ARPE-19 cells were cultured in exosome-depleted medium. sEVs were isolated from cell culture supernatants. A) Schematic representation of LAMP2 genomic sequence. There are 3 isoforms of LAMP2 (A, B, C) originated by the alternative splicing of exon 9. In red is the exon for the LAMP2A isoform. B) TEM images show isolated exosomes. Graphs show particle number and size using nanoparticle tracking system (NanoSight). LAMP2A KO maintains particle size and number. C) Western blot of cells extracts and sEVs fractions blotted with antibodies raised against the exosomal positive markers (CD63, FLOT1 and HSC70), exosome negative markers (CANX and ATP1A1) and KFERQ-containing proteins. LAMP2A KO leads to a decrease in the proteins levels of EEA1, EXOc2 and Histone3, which contain at least one KFERQ-motif each, as well as HSC70. D) Raw MSMS fragmentation spectrum of the peptide HHHAGYEQF, exclusively from LAMP2 isoform A, present only in WT samples. E). Schematic representation shows the rules for the identification of putative KFERQ-motifs. Graph shows the percentage of proteins with at least one KFERQ-motif in their sequence that are present in sEVs. F) Heatmaps of dowregulated proteins in sEVs after LAMP2A KO. 67% of Down-regulated proteins contain KFERQ-motifs. All samples were analyzed under the same experimental conditions. The results represent the mean ±SD of at least N=3 independent experiments (n.s. nonsignificant; *p < 0.05; **p < 0.01; ***p < 0.001).

EVs from the media supernatants of wild-type (WT) and LAMP2A KO ARPE-19 cells were isolated for MS-based proteome analysis. According to EVs guidelines^16^ we were able to isolate vesicles smaller than 200nm (that include exosomes) hereafter referred as sEVs. We confirmed that the size of the secreted sEVs of WT and LAMP2A KO cells is within the range of exosomes (∼100nm) by Transmission Electron Microscopy (TEM) as well as by laser scattering using NanoSight-tracking system (Fig.1B). TEM of ARPE-19 cells further shows that MVBs morphology is maintained after LAMP2A KO (SupFig.1B). Moreover we confirmed the presence of exosome positive markers CD63, Flotillin-1 and HSC70 and the absence of the exosome negative markers Calnexin and Na^+^K^+^ ATPase in our sEVs fractions (Fig.1C)^16, 17^. LC-MS/MS of isolated sEVs confirmed the presence of the peptide fragment HHHAGYEQF, specific for LAMP2A, only in WT sEVs (Fig.1D).

With the LC-MS/MS analysis of sEVs fractions we identified 2897 proteins (TableS1). Accordingly, the identified proteins are enriched in sEVs markers (SupFig.1C). Because LAMP2A is reported to bind to proteins that contain KFERQ-like motifs^15^, next we searched the identified proteins for these amino acid sequences. KFERQ-motifs are degenerated pentapeptide sequences recognized by the Heat shock 70 kDa protein 8 (HSC70) (Fig.1E). Initial estimates of the abundance of KFERQ-motifs in the proteome indicated that there are ∼25% of proteins containing such active/exposed motifs^18^. In our isolated sEVs, approximately 62% of all proteins contain a putative KFERQ motif (Fig.1E, TableS2), which represents a 2,5-fold enrichment. Further analysis of sEVs secreted from WT and LAMP2A KO cells shows that 303 proteins where significantly downregulated (p value <0,05) in LAMP2A KO sEVs, and that 203 of the downregulated proteins (67%) contain a putative KFERQ-motif (Fig.1F). Western Blot (WB) of LAMP2A KO sEVs shows a reduction in HSC70 levels, while confirming a decrease in the levels of randomly selected proteins from the LC-MS/MS data (Fig 1C). These findings suggest that LAMP2A KO leads to a decrease in the levels of proteins containing KFERQ-like motifs in sEVs.

### Loading of KFERQ-like motif tagged proteins in small extracellular vesicles is dependent on LAMP2A and HSC70

Based on previous reports^19, 20^, to address the mechanisms involved in the targeting of proteins containing KFERQ-motifs to sEVs, we used chimeric proteins consisting of a fluorescent protein, such as mCherry, and the KFERQ-motifs of a-synuclein (VKKDQ) and RNase A (KFERQ) separated by a peptide spacer (Fig.2A). WB of sEVs fractions from cells expressing either mCherry alone or mCherry fused to the targeting peptide, hereafter referred to as ExoSignal, showed that the ExoSignal tag increases mCherry presence in sEVs by ∼7 fold (Fig.2B,D). On the other hand, KO of LAMP2A from cells expressing mCherry-ExoSignal prevents the loading of the chimeric protein into sEVs (Fig.2C,D). Additionally, the rescue of LAMP2A expression is sufficient to restore the levels of mCherry-ExoSignal in sEVs (Fig.2E). mCherry-ExoSignal is present in sEVs fractions, positive for CD63 and with typical exosomal densities (SupFig.2A). Moreover, incubation of sEVs with trypsin showed that mCherry is resistant to degradation, while the sEVs membrane protein Cx43^6^ is not, indicating that mCherry-ExoSignal is incorporated into the lumen of sEVs (SupFig2.B). To further confirm that mCherry-ExoSignal is indeed loaded into exosomes, we down-regulated RAB27 thus inhibiting the release of exosomes^17, 21^. Rab27 depletion decreased sEVs secretion as well as the levels of mCherry-ExoSignal present in the sEVs fractions (Fig.2F, SupFig2.C).

**Figure 2.**
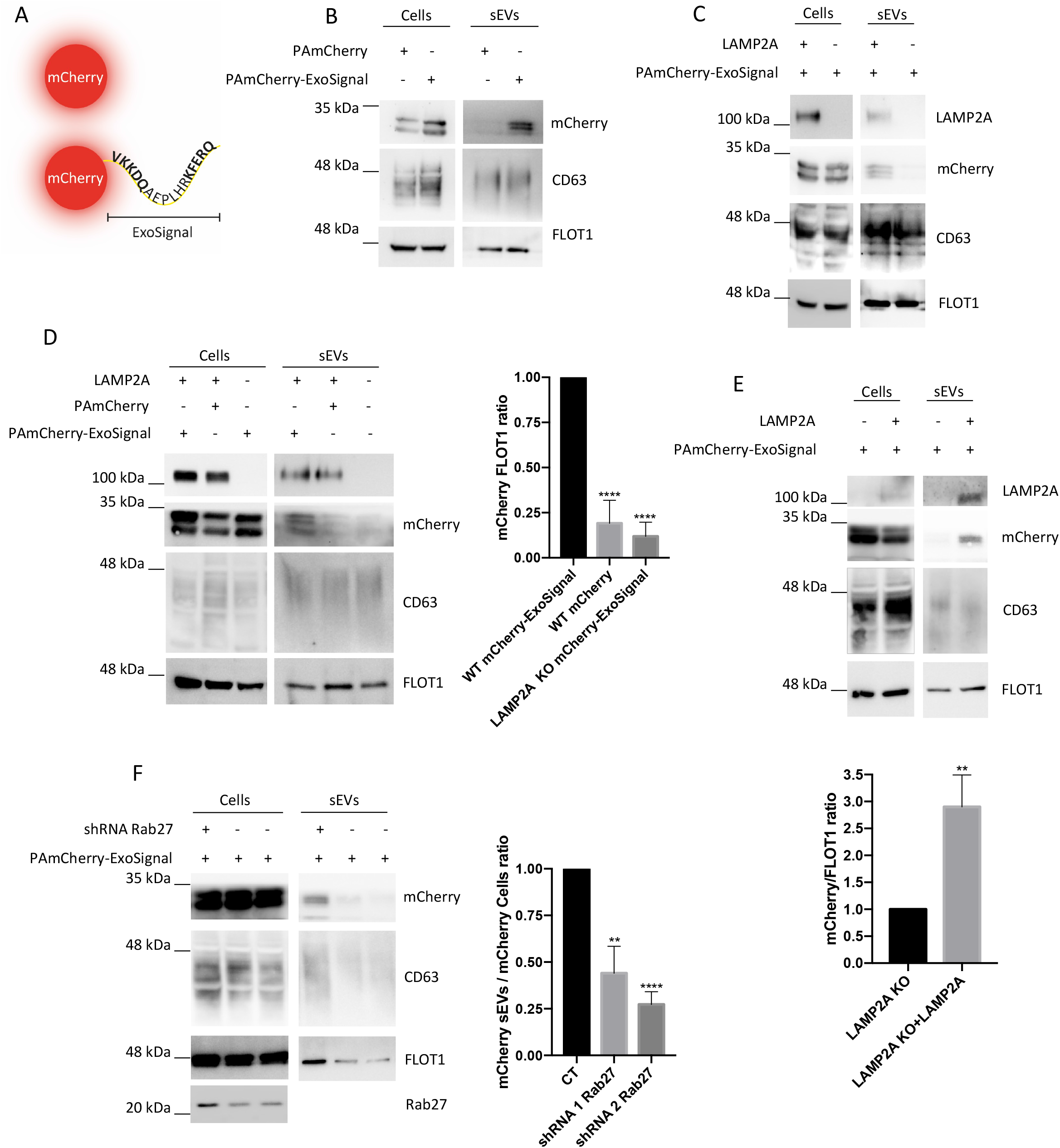
mCherry is presence in sEVs depends both on the ExoSignal and LAMP2A. WT and LAMP2A KO cells were transduced with lentiviral particles for the expression of either mCherry or mCherry-Exosignal. sEVs were isolated from cell culture supernatants. Schematic representation of mCherry tagged with the ExoSignal sequence. B,C,D) Western blot of cells extracts and isolated sEVs blotted with antibodies raised against CD63, FLOT1, LAMP2A and mCherry. The addition of the ExoSignal increases the levels of mCherry in sEVs. C, D) KO of LAMP2A decreases the levels mCherry-ExoSignal into exosomes. E) ARPE-19 cells LAMP2A KO were transduced with lentiviral particles to express human LAMP2A. Results show that rescuing LAMP2A expression is sufficient rescue the presence of mCherry-ExoSignal in sEVs. F) ARPE-19 cells expressing mCherry-Exosignal were transduced with adenoviral particles to express shRNA sequences for the depletion of Rab27. The release of sEVs loaded with mCherry-ExoSignal is decreased by the depletion of Rab27. All samples were analyzed under the same experimental conditions. The results represent the mean ±SD of at least N=3 independent experiments (n.s. nonsignificant; *p < 0.05; **p < 0.01; ***p < 0.001).

We hypothesize that HSC70 is involved in the targeting of proteins into nascent sEVs by binding and delivering proteins containing KFERQ-like motifs to endosomes. To assess if HSC70 has a role in loading of proteins into sEVs we used Pifithrin-μ (Pifi)^22^, a compound that acts by inhibiting the interaction of the HSC70 with its substrates. Immunoprecipitation data shows that incubation of cells with 7,5 μM of Pifi leads to an inhibition of the interaction between HSC70 and both mCherry-ExoSignal and LAMP2A (Fig.3A) and to a decrease mCherry-ExoSignal protein levels in isolated sEVs (Fig.3B). By immunoprecipitating mCherry from cells or sEVs we confirmed that, when tagged with the ExoSignal, both HSC70 and LAMP2A co-precipitate with mCherry (Fig.3C,D).

**Figure 3.**
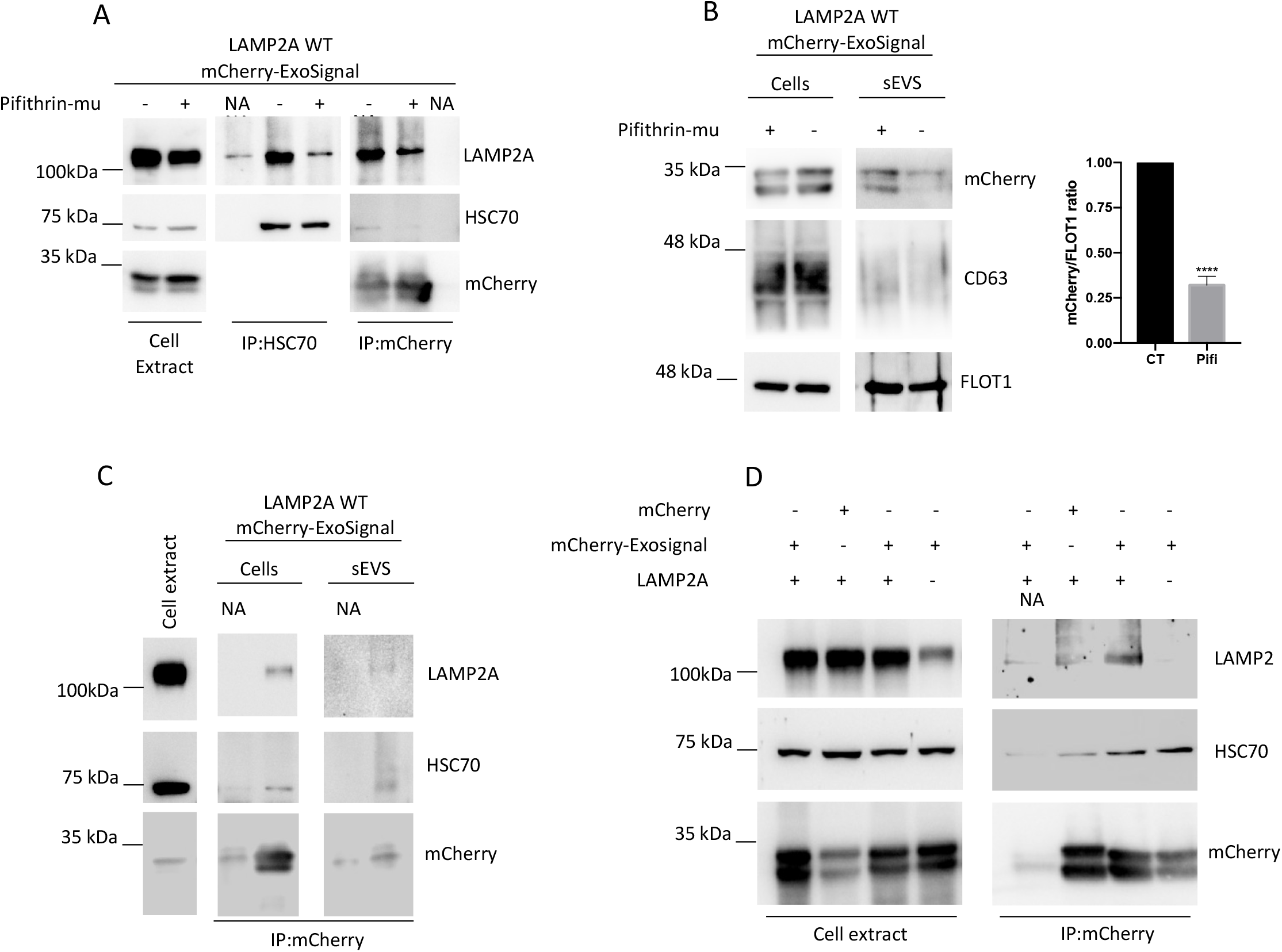
HSC70 is necessary for the presence of mCherry-ExoSignal in sEVs. WT and LAMP2A KO ARPE-19 cells were transduced using lentiviral particles containing vectors for the expression of either mCherry or mCherry-ExoSignal. A) Cells were maintained in the presence or absence or 7,5 μM of Pifithrin-mu (Pifi) for 12h. Immunoprecipitation experiments using antibodies raised against HSC70 and mCherry show that Pifi inhibits the interaction between HSC70 with PAmCherry-ExoSignal while also decreasing the interaction with LAMP2A. B) Cells were maintained in the presence or absence or 7,5 μM of Pifithrin-mu for 12h in Exosome-depleted medium. Western Blot of exosomal fractions using antibodies raised against CD63, FLOT1 and mCherry show that Pifi induces a decrease in the levels of mCherry-ExoSignal in sEVs. C) Cells were maintained in the exosome-depleted medium for 48h and exosomes were isolated from the media supernatant. Immunoprecipitation using antibodies raised against mCherry in exosomal samples show that PAmCherry-ExoSignal interacts with HSC70 and LAMP2A in sEVs. D) Immunoprecipitation in cell extracts using antibodies raised against mCherry show that mCherry-ExoSignal, but not mCherry alone, co-precipitates with HSC70 and LAMP2A. All samples were analyzed under the same experimental conditions. The results represent the mean ±SD of at least N=3 independent experiments (n.s. nonsignificant; *p < 0.05; **p < 0.01; ***p < 0.001).

Overall, data suggests that cytosolic proteins containing KFERQ-motifs are loaded into sEVs in a mechanism that involves both LAMP2A and HSC70.

### Proteins tagged with KFERQ-like motifs are loaded into endosomes early in the endocytic pathway

Fluorescent dextrans are commonly used to track fluid-phase endocytosis. When endocytosed, these compounds follow the endocytic pathway until they reach the lysosome^23^. We pulsed cells for 4h with Alexa647 dextran and chased its internalization for 20h to label lysosomes (Fig.4A) and used a 10 min pulse followed by a 5-10 min chase with Alexa488 dextran to identify Early Endosomes (EE), or a 15min chase for Late Endosomes (LE)^23^ (Fig.4A). For the imaging we used a photoactivable version of mCherry (PAmCherry). Our results (Fig.4A, SupFig.3A) show that while the majority of the intracellular PAmCherry-ExoSignal puncta is localized in lysosomal compartments, in images acquired as soon as 10 minutes after the chase, PAmCherry-ExoSignal is also present in Alexa488 dextran positive vesicles, indicating that is likely to be present at EE (Fig.4A). Additionally, LAMP2A KO cells show residual PAmCherry puncta (Fig4.A). After 15 min of chase, PAmCherry-ExoSignal puncta is in vesicles positive for both dextrans (SupFig.3A).

**Figure 4.**
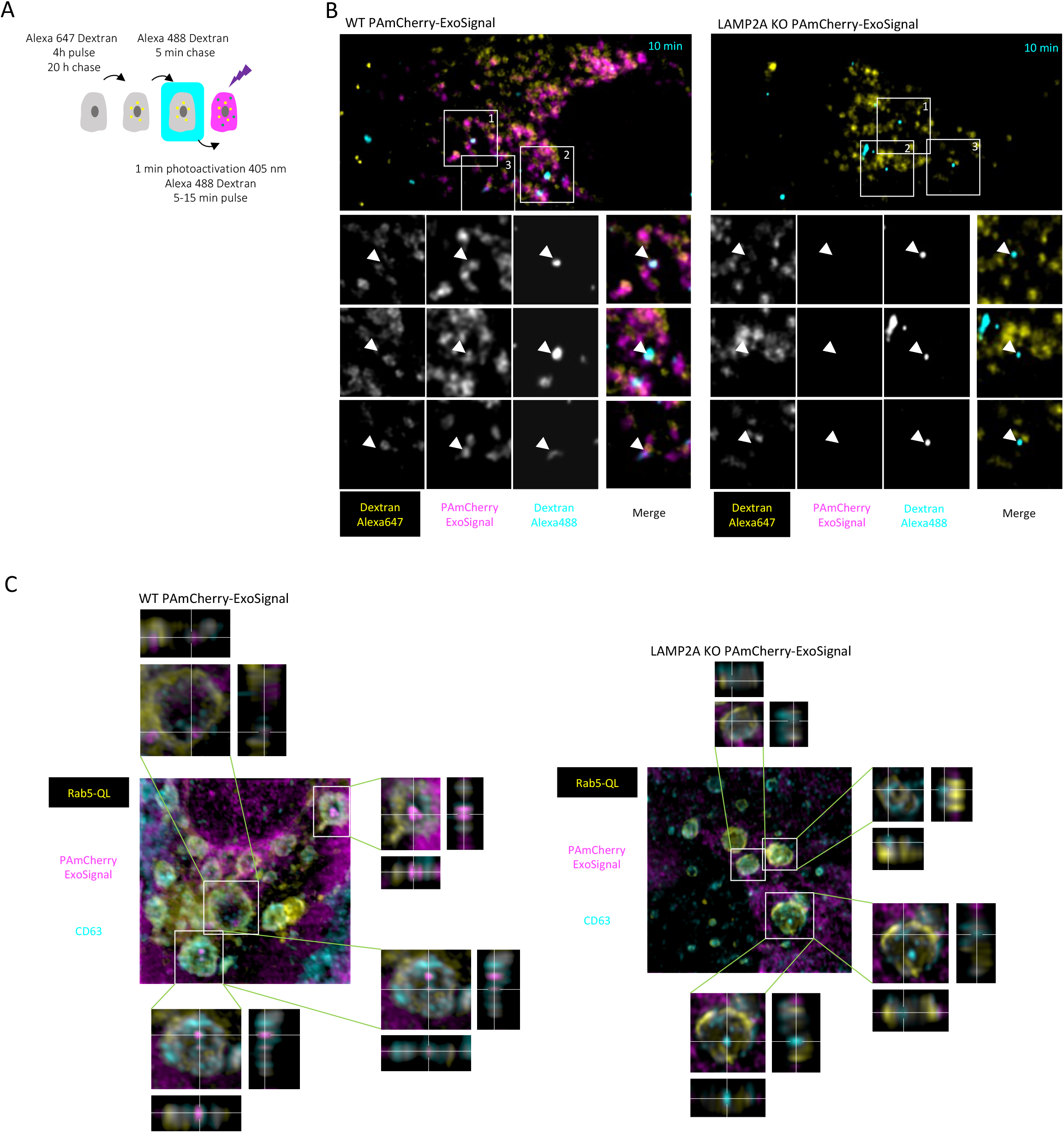
The PAmCherry-ExoSignal chimeric protein is sorted into early endocytic vesicles. WT and KO LAMP2A ARPE-19 cells were transduced with lentiviral particles containing vectors for the expression of PAmCherry-ExoSignal. A) Schematic representation of the dextran assay experimental setup. B) PAmCherry-ExoSignal was photoactivated with 405 nm laser for 1 min. Live cells were imaged in a Zeiss LSM 710 confocal microscope. Lysosomes are labelled in yellow (Alexa 647 dextran), endosomes in cyan (Alexa 488 dextran) and PAmCherry-ExoSignal in magenta. In WT cells, arrows at the 10 min time point show that some Alexa 488 dextran positive compartments, likely endosomes, are positive for PAmCherry-ExoSignal. In LAMP2A KO cells PAmCherry-ExoSignal puncta in residual. C) Cells were transfected with Rab5QL-GFP and incubated for 30 min with antibody against CD63 extracellular loop for 30min before fixation. Immunofluorescence using confocal microscopy shows that PAmCherry-ExoSignal puncta localizes in Rab5QL-GFP compartments, inside CD63 positive ILVs when LAMP2A is present. 3D images were reconstructed using Imaris software.

To further investigate the presence of ExoSignal tagged proteins in endosomes we used the constitutively active mutant of Rab5 (Q79L), that blocks the conversion EEs into LEs, resulting in the formation of very large hybrid endosomes that accumulate large ILVs^24^. Cells were incubated anti-CD63 antibody for 30 minutes before fixation^25^ to exclusively label CD63 within the endocytic pathway. Results show that mCherry-ExoSignal localizes in Rab5QL-GFP compartments, inside ILVs decorated with CD63 (Fig.4B). LAMP2A KO or the absence of the ExoSignal inhibits the targeting of the PAmCherry into CD63 positive ILVs (Fig.4B and SupFig.3B), while the rescue of LAMP2A expression is sufficient to restore the presence of PAmCherry-ExoSignal to ILVs (SupFig.3B).

Subsequently, we isolated endosomal compartments using a sucrose discontinuous gradient, according to de Araújo *et al.*^26^. We obtained 2 fractions, one enriched in the EE marker EEA1 and another enriched in the LE Rab7 (Fig.5A). The mature form of Cathepsin B (a lysosomal marker) is absent from both endosomal fractions. Data shows that mCherry-ExoSignal, but not untagged mCherry, is present both in early and late endosomal compartments, only when LAMP2A is present (Fig.5A).

**Figure 5.**
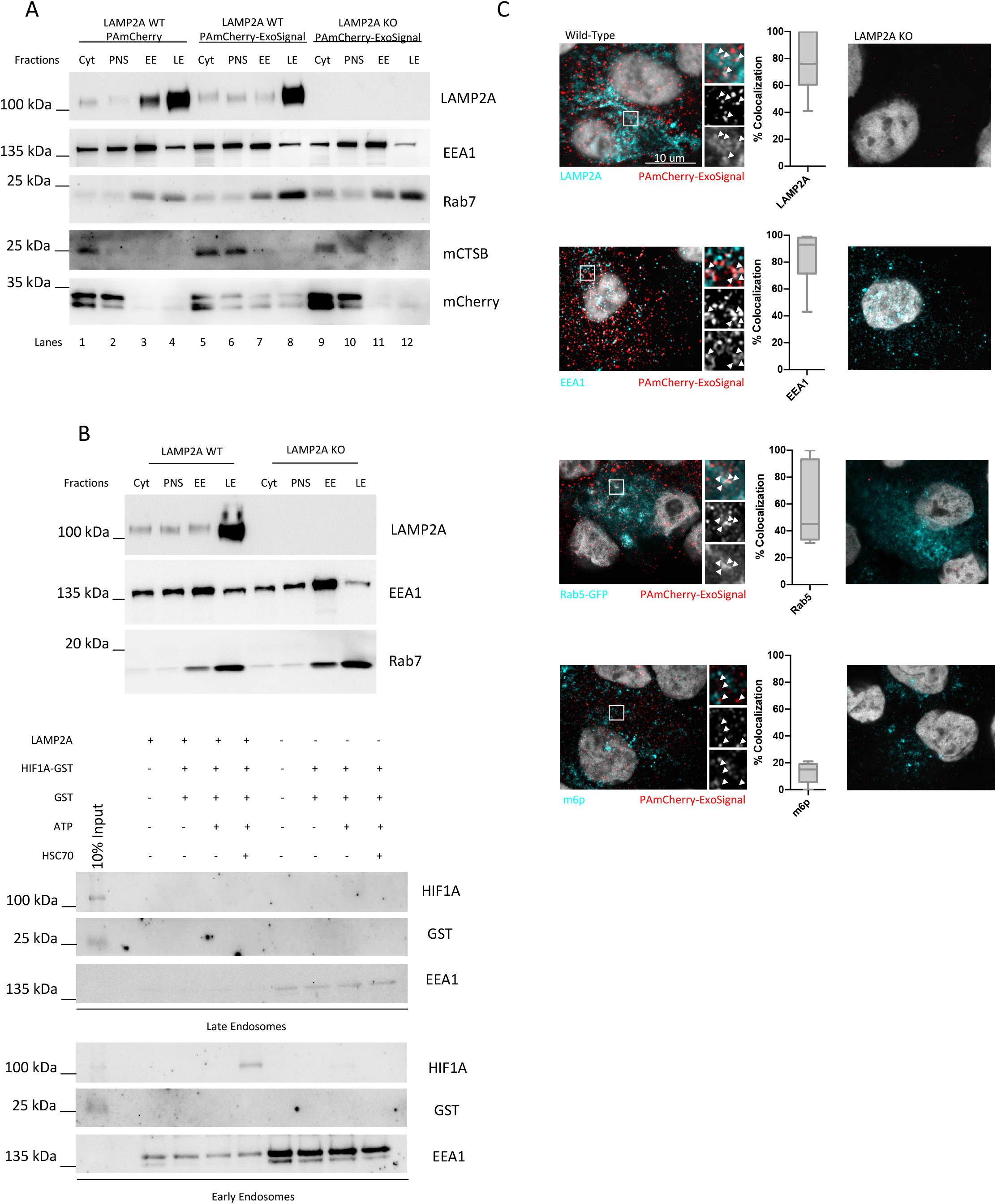
KFERQ-containing proteins are loaded into endosomes. Sucrose discontinuous gradients were performed as described in the protocol. Cytoplasmic (Cyt), Postnuclear supernatant (PNS), EEs and LEs enriched fractions of WT and KO LAMP2A cells were loaded into an SDS-PAGE. Western blots of isolated fractions were blotted with antibodies raised against the EE marker EEA1, the LE marker Rab7, the lysosome marker the Cathepsin B (CTSB, mature form), LAMP2A and mCherry. A) PAmCherry-ExoSignal, but not PAmCherry is present at both EE and LE compartments (compare lanes 3 and 4 with lanes 7 and 8, only when LAMP2A is present ) (compare lanes 7 and 8 with 11 and 12). B) Top panel depicts the separation of EEs and LEs. In the lower panel HIF1A-GST was incubated with freshly isolated vesicles in the presence/absence of ATP, the molecular chaperone HSC70, and LAMP2A. Samples were treated with trypsin to degrade all the protein that was not protected by the endosomal membrane. Western blot using antibodies raised against the GST-tag show that HIF1A-GST is translocated into EEs, in a LAMP2A and HSC70 dependent manner. C) Immunofluorescence using LSM 980 Aryscan confocal microscopy of cells fixed in methanol with antibodies against mCherry, LAMP2A, EEA1 and m6p or expressing Rab5-GFP. PAmCherry-ExoSignal shows high co-localization coefficient with LAMP2A, EEA1 and RAB5-GFP. KO LAMP2A cells expressing PAmCherry-ExoSignal show residual or no puncta.

Next we used the endosomal fractions for *in vitro* uptake assays, with recombinant HIF1A as a model substrate (hypoxia-inducible transcription factor 1 alpha). We had previously demonstrated that HIF1A contains an active KFERQ-motif^15, 17^. Incubation of ARPE-19 cells with the hypoxia mimetic compound CoCl2 showed that HIF1A is present is sEVs only when cells express LAMP2A (SupFig.4A). Isolation of sEVs from cells KO for HIF1A and the ubiquitin ligase Von Hippel-Lindau (VHL) (769-P)^27^, and either rescuing HIF1A expression or expressing a KFERQ-mutant HIF1A (HIF1A^AA^), shows that only WTHIF1A is present in sEVs (SupFig.4B). Moreover, subcellular fractioning using an Optiprep linear gradient showed that endogenous HIF1A localizes to endosomal fractions and that LAMP2A KO decreases HIF1A presence in those fractions (SupFig.4C). We incubated endosomes with GST (no KFERQ motif) or HIF1A-GST, in the presence or absence of recombinant HSC70 and ATP, followed by trypsin treatment to degrade recombinant HIF1A protein unprotected by the endosomal membrane (Fig.5B). Results show that only EE fractions incubated with HSC70, ATP and containing LAMP2A protected HIF1A-GST from degradation by trypsin (Fig.5B).

Confocal microscopy of cells after methanol fixation, to eliminate cytoplasmic mCherry and better resolve its vesicular localization^28^, show a high mCherry-ExoSignal co-localization coefficient with LAMP2A (76.67%±21.55) (Fig.5C, SupFig.5A), as well as with early endosomal markers such as Rab5 (60.7%±29.58) and EEA1 (84%±21.55), but shows low co-localization with late the endosome marker Mannose 6-phosphate (M6P) receptor (12.8%±8.012) (Fig.5C, SupFig.5A).

Previous reports indicated that some proteins containing KFERQ-like motifs are degraded in endosomes, independently of LAMP2A, by endosomal Microautophagy (e-Mi)^29^. Incubation of 2 typical e-Mi substrates, namely GAPDH and Aldolase, with endosomes isolated from WT and LAMP2 KO cells shows that both recombinant proteins are resistant to trypsin digestion in EEs and LEs, independently of LAMP2A and HSC70 (SupFig.5B).

Overall, data indicates that LAMP2A, as well as HSC70, have a role in loading KFERQ-containing proteins into endosomal ILVs early in the endocytic pathway.

### LAMP2A-mediated protein targeting to endosomes is independent of ESCRT machinery

TSG101 and VPS4b are components of the ESCRT-I and ESCRT-III machinery that are involved in ILVs formation and exosome biogenesis^30^. Additionally, the mechanism of e-Mi is dependent on TSG101 and VPS4b^29^. By depleting TSG101 we were able to observe a decrease the levels of CD63 and Flotilin-1 in sEVs fractions, indicating an inhibition in sEVs release. However, the levels of mCherry-ExoSignal present in sEVs from control or TSG101 depleted cells did not change (Fig.6A). On the other hand, VPS4b depletion increased the levels of the CD63 and Flotillin-1 in sEVs (Fig.6B). This is consistent with previous reports showing that VPS4b depletion led to an increase in exosome release^30^. Accordingly, the levels of mCherry-ExoSignal present in sEVs after VPS4b depletion are also increased (Fig.6B).

**Figure 6.**
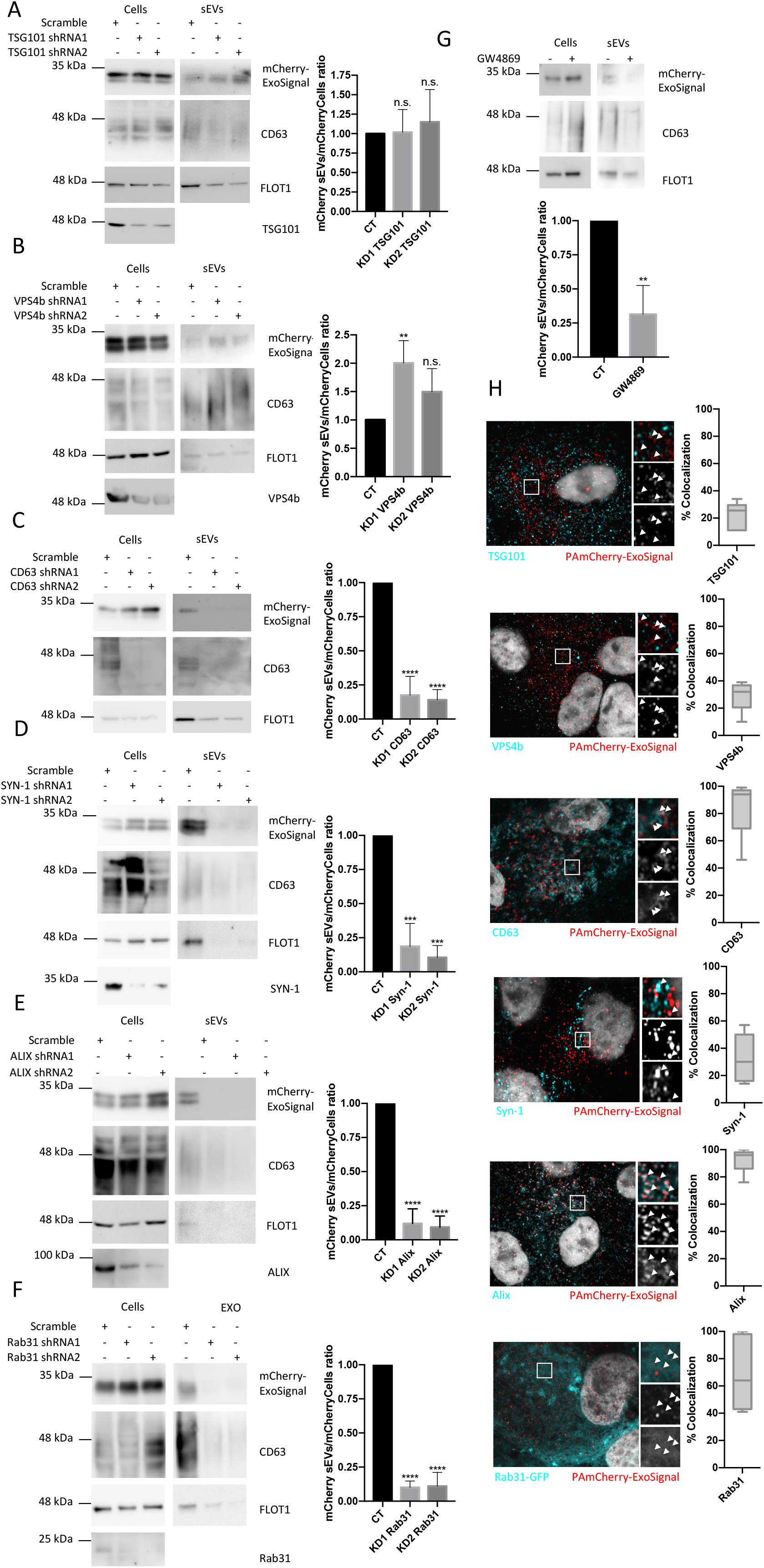
mCherry-ExoSignal is loaded into sEVs by ESCRT-independent mechanisms. WT ARPE-19 cells expressing mCherry-ExoSignal were transduced using lentiviral particles containing vectors for the expression of shRNAs. Cells were maintained in exosome-depleted medium for 48h. Western Blot of exosomal fractions using antibodies against CD63, FLOT1 and mCherry. A) TSG101 depletion decreases sEVs secretion but has no impact in the levels of PAmCherry-ExoSignal present in isolated sEVs. B) VPS4b depletion increases sEVs secretion as well as the levels of PAmCherry-ExoSignal present in isolated sEVs. C,D,E,F,G) CD63, Syn-1, Alix and Rab31 depletion, as well as ceramide synthesis inhibition by GW4869 decreased sEVs secretion and mCherry-Exosignal levels in the vesicles. H) Confocal images of methanol fixated cells show high colocalization coefficient of mCherry-ExoSignal with CD63, Alix and Rab31, and low colocalization coefficient for TSG101, VPS4b and Syn-1. All samples were analyzed under the same experimental conditions. The results represent the mean ±SD of at least N=3 independent experiments (n.s. nonsignificant; *p < 0.05; **p < 0.01; ***p < 0.001).

We next targeted ESCRT-independent mechanisms of exosome biogenesis^30^, including the sphingolipid ceramide^8^, the tetraspanine CD63^31^, the protein complex Syndecan-Syntenin-1-Alix^9^ and Rab31^32^. Both depletion of CD63, Syntenin-1, Alix and Rab31 using shRNA, or of ceramide levels using an inhibitor of ceramide synthesis (GW4869), inhibited sEVs secretion as well as mCherry-ExoSignal presence in the sEVs fraction (Fig.6C,D,E,F and G). Confocal microscopy images further show low co-localization between mCherry-ExoSignal with either TSG101 (22.5%±9.439) or VPS4b (28,83± 10.59) (Fig.6H). Conversely, images show high co-localization with CD63 (83.36%±18.96), Alix (92.33±8.093) and Rab31 (70.11%±26,49) while indicating lower than expected co-localization with Syntenin-1 (31,45±16,28) (Fig.6H).

Next, we immunoprecipitated LAMP2A or LAMP2B using antibodies raised for the cytoplasmic tail of each specific isoform. Results show substantial cross-reactivity between the so-called LAMP2B specific antibody and the LAMP2A protein isoform (Fig.7A). No such cross-reaction is apparent with the LAMP2A specific antibody (Fig.7A). Nonetheless, we were able to co-precipitate CD63 and ALIX, but not TSG101, after LAMP2A immunoprecipitation and *vice-versa* (Fig.7A,B). Conversely, in samples incubated with the less-specific anti-LAMP2B antibody, both CD63 and ALIX showed residual co-precipitation (Fig.7A).

**Figure 7.**
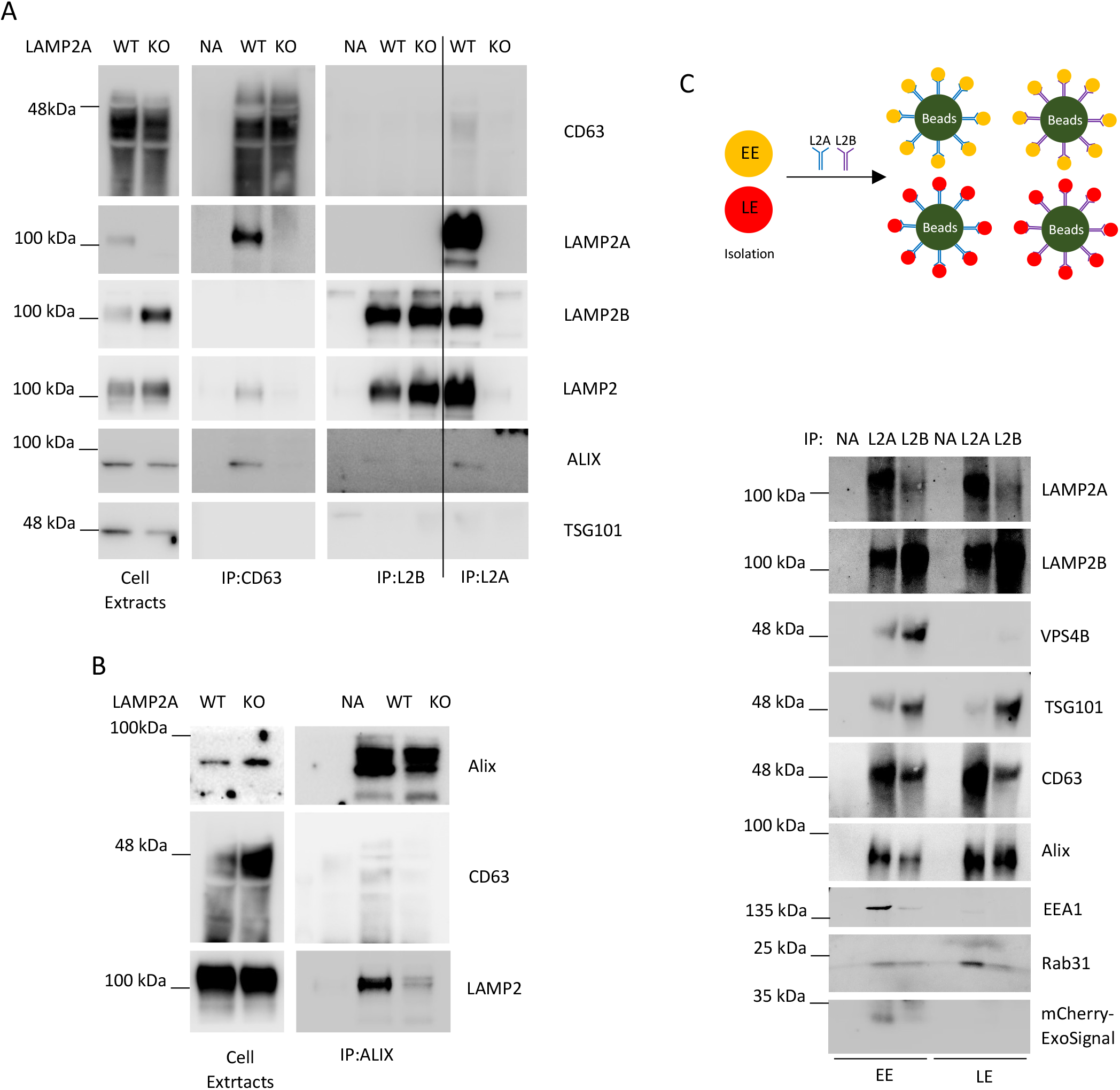
LAMP2A associates more closely to ESCRT-independent endosomal machinery. A,B) Immunoprecipitation experiments using antibodies raised against LAMP2, LAMP2A, LAMP2B, CD63, Alix and TSG101 show that LAMP2A, but not LAMP2B, co-precipitates with CD63 and Alix. TSG101 does not co-precipitate with either the LAMP2A or LAMP2B isoform. C) Top figure shows the schematic workflow of the experiment. EEs and LEs were further immunoprecipitated with antibodies raised against the LAMP2A or LAMP2B isoforms. WB show (bottom figure) VPS4B and TSG101 are enriched in LAMP2B pulled-down endosomes while CD63, Alix, RAB31 and mCherry-ExoSignal are enriched in LAMP2A enriched endosomes.

Subsequently we isolated endosomes and immunoprecipitated the subcellular compartments with either anti-LAMP2A or anti-LAMP2B antibodies. When comparing LAMP2A with LAMP2B endosomes the distribution of the different components is asymmetric. There are higher levels of TSG101 and VPS4b in endosome fractions enriched in LAMP2B and higher levels of CD63 and Rab31 in endosome fractions enriched in LAMP2A (Fig.7C). Alix is asymmetrically distributed towards LAMP2A endosomes only in EEs enriched fractions (Fig.7C). Accordingly, mCherry-ExoSignal levels are higher in EEs enriched in LAMP2A (Fig.7C).

Overall, evidences indicate that LAMP2A, but not LAMP2B, participates in the loading of proteins containing KFERQ-like motifs at the early endosomal membrane in association with CD63, Alix, Syntenin-1, Rab31 and ceramides.

### ExoSignal tagged proteins are sorted into circulating sEVs in zebrafish larvae

In a recent report we followed circulating exosomes in zebrafish larvae, as they travelled from one organ to another, using CD63 tagged with a fluorescent protein^33^. Building up from that concept we now created a new construct composed of untagged mCherry, followed by the self-cleaving Porcine Teschovirus-1 derived peptide (P2A)^34^ and GFP tagged with the ExoSignal (SupFig.7A). We injected the yolk syncytial layer (YSL) of zebrafish embryos, a highly secretory and irrigated organ of the zebrafish larvae^33^ and imaged the zebrafish caudal plexus, that seats immediately after the yolk sack (SupFig.7A). When compared with sham-injected larvae or untagged GFP, mCherry-P2A-GFP-ExoSignal injected larvae are positive for GFP in the caudal plexus, while mCherry signal is residual (Fig.8A, SupFig.7B; Smovie1,2,3,4,5). GFP-ExoSignal positive puncta, likely sEVs, circulate in the blood stream (Fig.8B, Smovie1,2,3 and 6 in higher magnification), and often stop at the vessel’s wall (Fig.8C, Smovie7). Moreover, sEVs isolated from zebrafish larvae^33^ contain GFP-ExoSignal, but not mCherry (SupFig.7C). Co-injection of a morpholino (MO) raised against zebrafish syntenin-a^33, 35^, the zebrafish homolog for human syntenin-1, to inhibit sEVs secretion (SupFig.7D), as well as a MO against zebrafish LAMP2A, decreased GFP-ExoSignal in the caudal plexus of zebrafish larvae (Fig.8A, Smovie 8 and 9).

**Figure 8.**
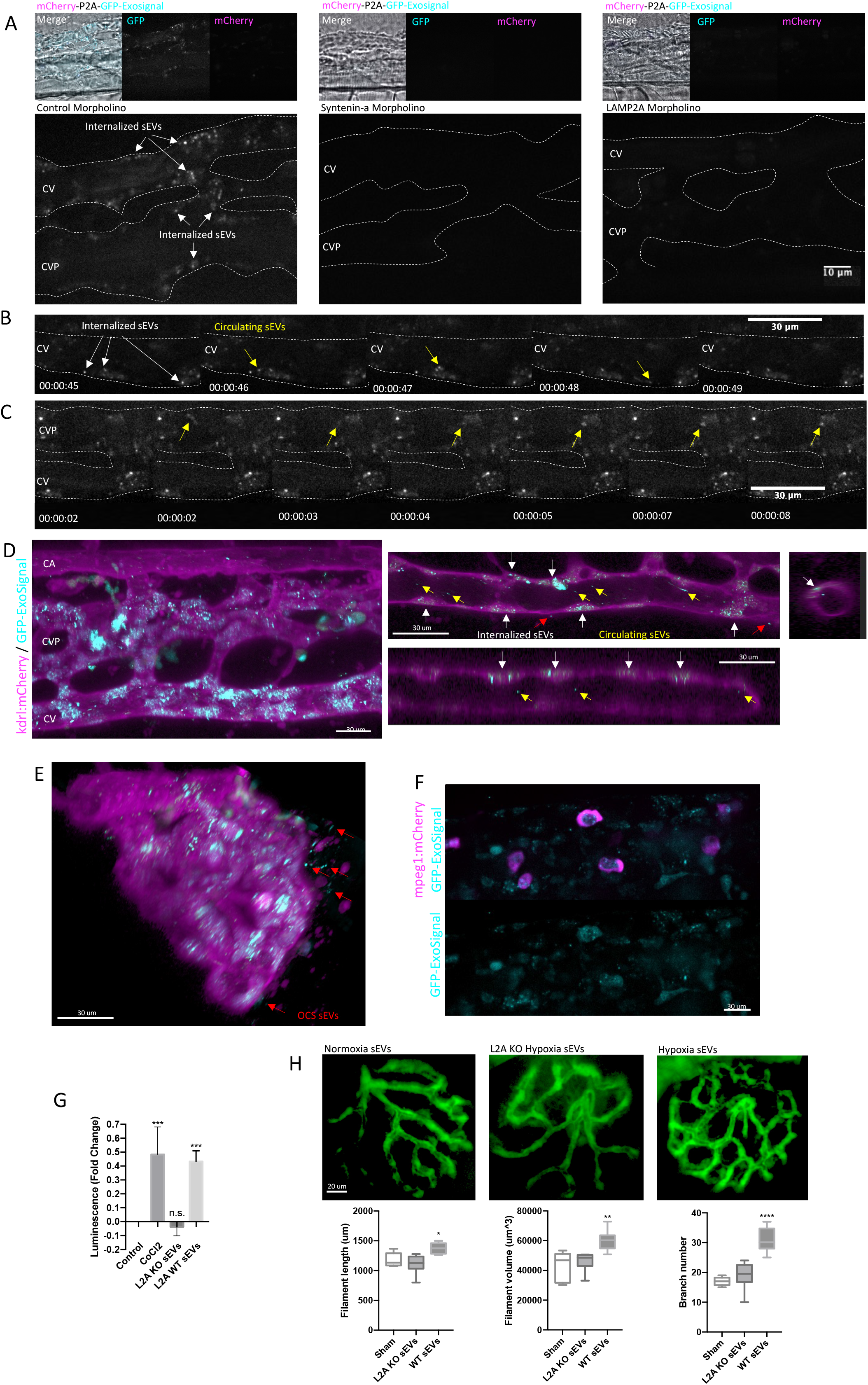
LAMP2A mediates loading of KFERQ-containing proteins into sEVs in the zebrafish larvae. Brightfield movies and still images of the zebrafish show the caudal artery (CA) and the caudal vein (CV) as well as a complex venous vascular network (CVP-caudal vascular plexus) located in the space between the vein and the artery. A) *casper* zebrafish at 1000 cell stage were injected at the YSL with a construct for the expression of GFP-ExoSignal-P2A-mCherry with either Control, Syntenin-a or LAMP2A Morpholinos. Confocal microscopy using a spinning disk show GFP-Exosignal is present in the zebrafish larvae caudal plexus at 3dpf. Syntenin-a or LAMP2A MO inhibit GFP-ExoSignal circulation in the caudal plexus. B,C) Time-lapse of Smovie 6 and 7 showing GFP-ExoSignal sEVs in the caudal plexus, with white arrows indicating incorporated sEVs and yellow arrows indicating circulating sEVs. D,E) 3D reconstruction and orthogonal view of *Tg(kdrl:mCherry)* larvae, expressing mCherry in endothelial cells, injected with GFP-ExoSignal construct. White arrows indicate internalized sEVs, yellow arrows indicate circulating sEVs and red arrows indicate outside circulatory network (OCV) sEVs. sEVs labeled with GFP-ExoSignal are present inside endothelial cells, in the vessel’s lumen as well as outside the OCV. F) *Tg(mpeg1:mCherry)* larvae, expressing mCherry in macrophages, injected with GFP-ExoSignal construct. GFP-Exosignal is present in macrophages located in the caudal plexus. G) HeLa cells expressing Luciferase under the control of a Hypoxia Response Element (HRE) were incubated with WT (loaded with HIF1A) and LAMP2A KO (missing HIF1A) sEVs. HIF1A from sEVs activates luciferase expression. H) WT and KO LAMP2A ARPE-19 cells were cultured in exosome-depleted medium incubated with 300μM of the hypoxia-mimetic agent CoCl_2_ for 12h. 0,25μg of isolated exosomes were injected in 3dpf zebrafish embryos expressing GFP in endothelial cells *Tg(fli1a:EGFP)*. At 5dpf, whole-mounted embryos where imaged. sEVs loaded with HIF1A are able to induce neovascularization in the zebrafish embryos. All samples were analyzed under the same experimental conditions. 3D reconstruction of the images and quantifications were performed using Imaris software. The results represent the mean ±SD of at least N=3 independent experiments (n.s. nonsignificant; *p < 0.05; **p < 0.01; ***p < 0.001).

3D reconstruction of the caudal plexus of *Tg(kdrl:mCherry)*, expressing mCherry in endothelial cells, shows that GFP-ExoSignal is present in the lumen of the blood vessels, as well as inside endothelial cells (Fig.8D), while no GFP signal is present in endothelial cells in the sham-injected or syntenin-a MO conditions (SupFig.7E). In some instances, GFP also localizes outside the vascular network (Fig.8E). By live imaging of *Tg(mpeg1:mCherry)*, expressing mCherry in macrophages, we can observe that macrophages moving within the caudal plexus have incorporated GFP-ExoSignal (Fig.8F, SupFig.7F, Smovie10 and 11).

Overall data shows that sEVs loaded with GFP-ExoSignal are released from the YSL into the circulation, travelling through the blood stream to reach the caudal plexus, where they are incorporated by endothelial cells and infiltrating macrophages.

### LAMP2A regulates intercellular transfer of hypoxia signalling through the HIF1A transcription factor

We incubated cells 769-P cells, KO for HIF1A^27^, with the sEVs loaded with HIF1A. Subcellular fractioning of 769-P cells shows that sEVs transported HIF1A reaches the cytosol as well as the nucleus (SupFig.7G). HeLa cells expressing an HIF1A reporter gene, with Luciferase under the control of a Hypoxia Response Element (HRE) consensus sequence and incubated with WT (loaded with HIF1A) and LAMP2A KO (lacking HIF1A) sEVs, shows that only sEVs secreted by LAMP2A WT cells induce luciferase expression (Fig.8G).

HIF1A adaptative response to hypoxia involves the formation of new blood vessels, or neovascularization^36^. By injecting a zebrafish transgenic line *Tg(fli1a:EGFP),* expressing GFP in endothelial cells, with sEVs from WT cells we were able to increase the number of vascular branches, including longer blood vessels with greater volume, in the retinal vascular network (Fig.8H), while sEVs isolated from LAMP2A KO cells failed to do so. Overall, data indicates that cells expressing LAMP2A release sEVs, carrying HIF1A, capable of reaching receiving cells. Once inside the cells HIF1A is transported to the nucleus of normoxic cells to activate HIF1A mediated gene transcription.

## Discussion

In this manuscript we propose a mechanism whereby LAMP2A, in coordination with HSC70 and additional molecular players, participates in the loading of cytosolic proteins into a subpopulation of exosomes. The targeted proteins contain amino acid sequences, biochemically related to the KFERQ-motif, a pentapeptide sequence previously involved in two selective forms of autophagy: Chaperone-Mediated Autophagy (CMA)^15, 37^ and endosomal Microautophagy (e-Mi)^29^. LAMP2A was initially thought to be involved exclusively in CMA, by mediating substrate translocation across the lysosomal membrane in a process that was dependent on the chaperone HSC70 ^15, 37^. We now suggest that LAMP2A also participates in the loading of proteins into ILVs at the early endosomal membrane. Our data also suggests that endosomal LAMP2A does not participate in the events leading to ILV formation by the inward budding of the endosomal limiting membrane. Instead, it is likely that by binding to HSC70 and associated KFERQ-containing proteins, LAMP2A captures its cargo during ILVs biogenesis.

Additionally, results show that the loading of proteins containing KFERQ-motifs into endosomes is likely to be restricted to EEs. This is not surprising since it is possible that the ability to generate ILVs is greater in EEs than in LEs. For example, it was shown that CD63 is ∼7x enriched in the ILVs of LEs as compared to the endosomal limiting membrane^38^. Nonetheless it is intriguing that the loading of the two e-MI substrates GAPDH and Aldolase ^29^, which were described to be independent of LAMP2A, occurs both in EEs and LEs enriched fractions. In this case, it is likely that the machinery associated to e-MI, including the ESCRT machinery^29^, is active in later endosomal compartments, while LAMP2A and other ESCRT-independent machinery is not, but further research is needed to clarify these discrepancies.

Our data indicates that LAMP2A-mediated loading of mCherry-ExoSignal into exosomes is ESCRT-independent, while involving CD63, ALIX, Syntenin-1, Rab31 and ceramides. Interestingly, after the pull-down of endosomes enriched in either the LAMP2A or LAMP2B isoforms, the different endosomal components were asymmetrically segregated, with the ESCRT machinery preferentially localizing to LAMP2B enriched endosomes while the other components segregated preferentially to LAMP2A enriched endosomes. These evidences support a model in which endosomal compartments within a cell may contain distinct molecular machineries, which will impact the final contents of ILVs and, as a consequence, of exosomes.

To assess the biological impact of LAMP2A-mediated exosomal loading we used the zebrafish model. Only when GFP was tagged with the ExoSignal were we able to observe exosomes being transferred from the YSL to endothelial cells and macrophages within the caudal plexus via circulation. Both depletion of exosome secretion by using the syntenin-a MO^33^ , or using a MO for LAMP2A, were able to inhibit the presence of circulating GFP-ExoSignal, further corroborating our previous *in vitro* observations.

Our lab had previously reported that the transcription factor HIF1A is a substrate for CMA^15, 39^. Data presented here shows that HIF1A is selectively targeted to exosomes, by the virtue of its KFERQ-motif and LAMP2A. Exosomal HIF1A can activate HIF1A signaling in normoxic cells both *in vitro* and in the zebrafish animal model, acting as second hand messenger of hypoxia.

In this study we propose that LAMP2A mediates the loading of proteins into exosomes, in a process that is ESCRT-independent and involves HSC70 recognition of active KFERQ-motifs. Endosomal proteins such as CD63, Syntenin-1, Alix, Rab31, as well as ceramides, appear to be required for the selective loading, although its role is not yet clear. Evidences also suggest that this mechanism may impact intercellular and interorgan communication by promoting the loading of the hypoxia transcription factor HIF1A into exosomes with the potential to activate hypoxia response in distant normoxic cells and tissues.

## Methods

### 1. Cell culture and treatments

The human cell lines ARPE-19 (of retinal pigment epithelium) and 769-P (from renal cell adenocarcinoma) were cultured in DMEM with Glutamine (Dulbecco’s modified Eagle’s medium) (Biowest) supplemented with 10% fetal bovine serum (FBS, Biowest), Penicillin/Streptomycin (100 U/ml:100 μg/ml, Gibco). Cells were cultured at 37°C under 5% CO_2_. When appropriate, cells were treated with the following agents: 300μM Cobalt Chloride (CoCl_2_, Sigma-Aldrich), 7.5μM Pifithrin-μ (Merk) and 50nM Bafilomycin A1 (Apollo Scientific)

### Antibodies and reagents

The following antibodies were used: mouse anti-LAMP2 clone H4B4, dilution of 1:1000 (Western Blot WB) 1:100 (Immunofluorescence IF) (Santa Cruz Biotechnology, SC756); rabbit anti-LAMP2A dilution of 1:500 (Western Blot WB), 1:100 (Immunofluorescence IF) (Abcam, ab18528); rabbit anti-LAMP2B dilution of 1:500 (Abcam, ab18529); mouse anti-actin, dilution of 1:2000 (Sigma, AS441); goat anti-HIF1A, dilution of 1:1000 (Sicgen, AB0112-200); goat anti-CD63 dilution 1:1000 (Sicgen, AB0047-200); mouse anti-CD63 dilution 1:100 (IF) (Santa Cruz Biotechnology, MX-49.129.5); rabbit anti-Flotillin-1, dilution of 1:500 (Santa Cruz Biotechnology, H-104); goat anti-CTSB/Cathepsin B clone S-12, dilution of 1:500 (Santa Cruz Biotechnology, SC-6493); mouse anti-Alix dilution of 1:500 (Santa Cruz Biotechnology, sc-53540); mouse anti-TSG101 dilution 1:500 (GeneTex, GTX70255); goat anti-GST dilution 1:500 (Sicgen, AB9919-500); goat anti-EEA1 dilution 1:500 (Sicgen, AB0006-200); goat anti-mCherry dilution 1:500 (WB) 1:100 (IF) (Sicgen, AB0040-200); rabbit anti-dsRed dilution 1:100 (IF) (Clontech, 632496); mouse anti-M6PR dilution 1:100 (IF) (Abcam, ab2733); rat anti-HSC70 clone 1B5, dilution of 1:1000 (Stressgen, ADI-SPA-815); goat anti-Rab27 dilution 1:500 (Sicgen, AB7223-200); goat anti-Tubulin dilution 1:2000 (Sicgen, AB0046-200); mouse Lamin B1 dilution 1:500 (Santa Cruz Biotechnology, sc-374015); goat anti-Calnexin dilution 1:1000 (Sicgen, AB0041-200); goat anti-Na^+^/K^+^ATPase dilution 1:1000 (Sicgen, AB0306-200); goat anti-GAPDH dilution 1:2000 (Sicgen, AB0049-200); mouse anti-ExoC2 dilution 1:500 (Novus Biological, NBP1-83786); rabbit anti-Histone 3 1:1000 (Sigma, H0164); anti-Vps4b 1:500 (SCBT, sc-133122), goat anti-GFP 1:1000 (Sicgen, AB0020-200), goat anti-Rab31 1:500 (Sicgen, AB0068-200); mouse anti-syntenin-1 (SCBT, sc-515538) and horseradish peroxidase-conjugated secondary goat anti-mouse (Bio-Rad, 170-6516), goat anti-rabbit (Bio-Rad, 170-6515), rabbit anti-goat (Bio-Rad, 172-1034) and goat anti-rat (Invitrogen, A10549) dilution of 1:5000. Alexa Fluor 568-conjugated donkey anti-goat (Invitrogen, A11057), Alexa Fluor 488-conjugated donkey anti-goat (Invitrogen, A32814), Alexa Fluor 488-conjugated donkey anti-rabbit (Invitrogen, A32790), Alexa Fluor 568-conjugated donkey anti-goat (Invitrogen, A11057), Alexa Fluor 546-conjugated donkey anti-mouse (Invitrogen, A10036), donkey anti-rabbit Cy3 (Abcam, ab6566), donkey anti-mouse Cy5dilution of 1:250. Dextran, Alexa Fluor™ 488; 10,000 MW, Anionic, Fixable (Thermo Fisher Scientific, D22910). Dextran, Alexa Fluor™ 647; 10,000 MW, Anionic, Fixable (Thermo Fisher Scientific, D22914). DAPI (4’,6-Diamidino-2-Phenylindole, Dihydrochloride) (Invitrogen, D1306). Protein G–Sepharose (GE Healthcare, 17-0618-01). Dynabeads Protein G (Invitrogen, 10004B); Nitrocellulose membranes (GE Healthcare, 88018). ECL (GE Healthcare, RPN1235). Protease Inhibitor Cocktail (Sigma, P8340). Optiprep iodixanol density media (Sigma-Aldrich, D1556). ONE-Glo™ Luciferase Assay System (Promega, E6110), Pierce BCA Protein Assay Kit (Thermo Fisher Scientific, 23225).

### Peptide Sample Preparation for MS

The sEVs protein solution containing SDS and dithiothreitol (DTT) was loaded onto filtering columns and washed exhaustively with 8 M urea in HEPES buffer ^40^. Proteins were reduced with DTT and alkylated with IAA. Protein digestion was performed by overnight digestion with trypsin sequencing grade (Promega).

### Nano-LC-MS/MS Analysis

Peptide samples were analyzed by nano-LC-MSMS (DionexRSLCnano 3000) coupled to a Q-Exactive Orbitrap massspectrometer (Thermo Scientific). Briefly, the samples (5μl) were loaded onto a custom-made fused capillary precolumn (2 cm length, 360μm OD, 75μm ID) with a flow of 5μlper min for 7 min. Trapped peptides were separated on a custom-made fused capillary column (20 cm length, 360 μm outer diameter, 75 μm inner diameter) packed with ReproSil-PurC18 3-μm resin (Dr. Maish, Ammerbuch-Entringen, Germany) with a flow of 300 nl per minute using a linear gradient from 92% A (0.1% formic acid) to 28% B (0.1% formic acid in 100 acetonitrile) over 93 min followed by a linear gradient from 28 to 35% B over 20 min at a flow rate of 300 nl per minute. Mass spectra were acquired in positive ion mode applying automatic data-dependent switch between one Orbitrap survey. MS scan in the mass range of 400 to 1,200 m/z followed by higher-energy collisional dissociation (HCD) fragmentation and Orbitrap detection of the 15 most intense ions observed in the MS scan. Target value in the Orbitrap for MS scan was 1,000,000 ions at a resolution of 70,000 at m/z 200. Fragmentation in the HCD cell was performed at normalized collision energy of 31 eV. Ion selection threshold was set to 25,000 counts and maximum injection time was 100 ms for MS scans and 300 and 500 ms for MSMS scans. Selected sequenced ions were dynamically excluded for 45 s.

### MS Database Search

The obtained data from the 36 LC-MS runs were searched using VEMS ^41, 42^ and MaxQuant ^43^. A standard proteome database from UniProt (3AUP000005640), in which common contaminants were included, was also searched. Trypsin cleavage allowing a maximum of four missed cleavages was used. Carbamidomethyl cysteine was included as fixed modification. Methionine oxidation, N-terminal protein acetylation, lysine acetylation, lysine diglycine, and S, T, and Y phosphorylation were included as variable modifications; 10 ppm mass accuracy was specified for precursor ions and 0.01 m/z for fragment ions. The FDR for protein identification was set to 1% for peptide and protein identifications. No restriction was applied for minimal peptide length for VEMS search. Identified proteins were divided into evidence groups as defined^41^.

### MS Data analysis

Statistical and bioinformatics analysis of mass spectrometry data were performed in the statistical programming language R. Quantitative data from MaxQuant and VEMS were analyzed in R statistical programming language. IBAQ and protein spectral counts from the two programs were preprocessed by three approaches: 1) removing common MS contaminants followed by log2 (x + 1) transformation, 2) removing common MS contaminants followed by log2 (x + 1) transformation and quantile normalization, 3) removing common MS contaminants followed by log2 (x + 1) transformation, quantile normalization and abundance filtering to optimize overall Gaussian distribution of the quantitative values. For simplicity only the quantile normalized quantitative data are presented in the manuscript. Statistical differences were calculated by utilizing the R package limma ^44^. Correction for multiple testing was applied using the method of Benjamini & Hochberg ^45^. Comparison of EV markers and potential contaminants in current study with previous study was performed as previously described ^46^. Functional enrichment of KEGG, cellular component, molecular function and biological process were calculated by using the hypergeometric probability test as previously described ^47, 48^. For the search of KFERQ-like motifs we used the online tool KFERQ finder (https://rshine.einsteinmed.org/).

### Plasmids for transfection

For this work we used the following plasmids: pT81 HRE (x3)-Luciferase ^49^ and pEGFP-Rab5QL.

### CRISPR-Cas9

ARPE-19 cells were transduced with the lentiviral vector pCW-Cas9 (a gift from Eric Lander & David Sabatini, Addgene plasmid # 50661; http://n2t.net/addgene:50661; RRID:Addgene_50661) ^50^ for Doxycycline-induced expression of SpCas9. Infected cells were selected with puromycin. Cells were allowed to grow in very low confluency and a one cell colony was selected and allowed to reach confluency. The selected isotype was used from this point on. The gRNA targeting LAMP2A (5’-AGTACTTATTCTAGTGTTGC-3’) was expressed under U6 promoter. The expression cassette containing U6 promoter, target sequence, PAM sequence, gRNA and the termination signal for Pol III. This sequence was flanked by attB1/attB2, was synthesized by GeneCust and subcloned into pLenti6, containing blasticidin resistance (Thermo Scientific), using a Gateway adapted vector using BP/LR Clonase II (Thermo Scientific), according to the manufacturer’s instructions. The guide was designed using the online Guide Design Resources tool, from Zhang Lab., (https://zlab.bio/guide-design-resources).

For the KO of LAMP2A isoform cells were incubated with Doxycycline for 48h and were subsequently allowed to grow in very low confluency. Individual colonies were allowed to reach confluency and the KO of LAMP2A was confirmed by DNA sequencing and Western Blot.

### Lentiviral plasmids for protein expression

All lentiviral particles were produced by co-transfection of pLenti6 and pMD 2.G (VSV-G protein) and psPAX2 (Rev and Pol proteins) into a producer cell line 293STAR RDPro (ATCC). Recombinant viral particles were harvested 48 days later, cleared for cell debris by centrifugation at 3200 g for 10 min and used. pLenti6-wtHIF1A and KFERQ-mutated pLenti6-AAHIF1A ^15^ were obtained from the Molecular Biology Platform (CEDOC, Lisbon). PA-mCherry and PA-mCherry ExoSignal were flanked by attB1/attB2 Gateway (ThermoScientific) with the sequences: (5’ATGGTGAGCAAGGGCGAGGAGGATAACATGGCCATCATTAAGGAGTTCATGCGCTTCAAGGTGCACATGGAGGGGTCCGTGAACGGCCACGTGTTCGAGATCGAGGGCGAGGGCGAGGGCCGCCCCTACGAGGGCACCCAGACCGCCAAGCTGAAGGTGACCAAGGGCGGCCCCCTGCCCTTCACCTGGGACATCCTGAGCCCTCAGTTCATGTACGGCTCCAATGCCTACGTGAAGCACCCCGCCGACATCCCCGACTACTTTAAGCTGTCCTTCCCCGAGGGCTTCAAGTGGGAGCGCGTGATGAAATTCGAGGACGGCGGCGTGGTGACCGTGACCCAGGACTCCTCCCTGCAGGACGGCGAGTTCATCTACAAGGTGAAGCTGCGCGGCACCAACTTCCCCTCCGACGGCCCCGTGATGCAGAAGAAGACCATGGGCTGGGAGGCCCTCTCCGAGCGGATGTACCCCGAGGACGGCGCCCTGAAGGGCGAGGTCAAGCCCAGAGTGAAGCTGAAGGACGGCGGCCACTACGACGCTGAGGTCAAGACCACCTACAAGGCCAAGAAGCCCGTGCAGCTGCCCGGCGCCTACAACGTCAACCGCAAGCTGGACAT CACCAGCCACAACGAGGACTACACCATCGTGGAGCAGTACGAGAGAGCCGAGGGCCGCCACTCCACCGGCGGCATGGACGAGCTGTACAAG-3’)

The sequences were synthesized and cloned into pUC57 (GeneCust). PA-mCherry and PA-mCherry ExoSignal were subcloned into pLenti6 (Thermo Scientific) using a Gateway adapted vector using BP/LR Clonase II (Thermo Scientific) according to the manufacturer’s instructions.

Human LAMP2A was synthesized into pUC57 Plasmid with the sequence: (5’-CAAGTTTGTACAAAAAAGCAGGCTCTCGAGCAatgGTGTGCTTCCGCCTCTTCCCGGTTCCGGGCTCAGGGCTCGTTCTGGTCTGCCTAGTCCTGGGAGCTGTGCGGTCTTATGCATTGGAACTTAATTTGACAGATTCAGAAAATGCCACTTGCCTTTATGCAAAATGGCAGATGAATTTCACAGTACGCTATGAAACTACAAATAAAACTTATAAAACTGTAACCATTTCAGACCATGGCACTGTGACATATAATGGAAGCATTTGTGGGGATGATCAGAATGGTCCCAAAATAGCAGTGCAGTTCGGACCTGGCTTTTCCTGGATTGCGAATTTTACCAAGGCAGCATCTACTTATTCAATTGACAGCGTCTCATTTTCCTACAACACTGGTGATAACACAACATTTCCTGATGCTGAAGATAAAGGAATTCTTACTGTTGATGAACTTTTGGCCATCAGAATTCCATTGAATGACCTTTTTAGATGCAATAGTTTATCAACTTTGGAAAAGAATGATGTTGTCCAACACTACTGGGATGTTCTTGTACAAGCTTTTGTCCAAAATGGCACAGTGAGCACAAATGAGTTCCTGTGTGATAAAGACAAAACTTCAACAGTGGCACCCACCATACACACCACTGTGCCATCTCCTACTACAACACCTACTCCAAAGGAAAAACCAGAAGCTGGAACCTATTCAGTTAATAATGGCAATGATACTTGTCTGCTGGCTACCATGGGGCTGCAGCTGAACATCACTCAGGATAAGGTTGCTTCAGTTATTAACATCAACCCCAATACAACTCACTCCACAGGCAGCTGCCGTTCTCACACTGCTCTACTTAGACTCAATAGCAGCACCATTAAGTATCTAGACTTTGTCTTTGCTGTGAAAAATGAAAACCGATTTTATCTGAAGGAAGTGAACATCAGCATGTATTTGGTTAATGGCTCCGTTTTCAGCATTGCAAATAACAATCTCAGCTACTGGGATGCCCCCCTGGGAAGTTCTTATATGTGCAACAAAGAGCAGACTGTTTCAGTGTCTGGAGCATTTCAGATAAATACCTTTGATCTAAGGGTTCAGCCTTTCAATGTGACACAAGGAAAGTATTCTACAGCTCAAGACTGCAGTGCAGATGACGACAACTTCCTTGTGCCCATAGCGGTGGGAGCTGCCTTGGCAGGAGTACTTATTCTAGTGTTGCTGGCTTATTTTATTGGTCTCAAGCACCATCATGCTGGATATGAGCAATTTTAGGGTACCACCCAGCTTTCTTGTACAAAGTGGACGCGT-3’) Human LAMP2A was subcloned into pLenti6 (Thermo Scientific), a Gateway adapted vector using BP/LR Clonase II (Thermo Scientific), according to the manufacturer’s instructions.

P2A-GFP-ExoSignal was synthesized and subcloned into the p3E plasmid with the sequence. (5’-TACAGGTCACTAATACCATCTAAGTAGTTGATTCATAGTGACTGCATATGTTGTGTTTTACAGTATTATGTAGTCTGTTTTTTATGCAAAATCTAATTTAATATATTGATATTTATATCATTTTACGTTTCTCGTTCAACTTTCTTGTACAAAGTGGAGATCTatgGGAAGCGGACGTACTAACTTCAGCCTGCTGAAGCAGGCTGGAGACGTGGAGGAGAACCCTGGACCTGCTCGAGCAGTGAGCAAGGGCGAGGAGCTGTTCACCGGGGTGGTGCCCATCCTGGTCGAGCTGGACGGCGACGTAAACGGCCACAAGTTCAGCGTGTCCGGCGAGGGCGAGGGCGATGCCACCTACGGCAAGCTGACCCTGAAGTTCATCTGCACCACCGGCAAGCTGCCCGTGCCCTGGCCCACCCTCGTGACCACCCTGACCTACGGCGTGCAGTGCTTCAGCCGCTACCCCGACCACATGAAGCAGCACGACTTCTTCAAGTCCGCCATGCCCGAAGGCTACGTCCAGGAGCGCACCATCTTCTTCAAGGACGACGGCAACTACAAGACCCGCGCCGAGGTGAAGTTCGAGGGCGACACCCTGGTGAACCGCATCGAGCTGAAGGGCATCGACTTCAAGGAGGACGGCAACATCCTGGGGCACAAGCTGGAGTACAACTACAACAGCCACAACGTCTATATCATGGCCGACAAGCAGAAGAACGGCATCAAGGTGAACTTCAAGATCCGCCACAACAT CGAGGACGGCAGCGTGCAGCTCGCCGACCACTACCAGCAGAACACCCCCATCGGCGACGGCCCCGTGCTGCTGCCCGACAACCACTACCTGAGCACCCAGTCCGCCCTGAGCAAAGACCCCAACGAGAAGCGCGATCACATGGTCCTGCTGGAGTTCGTGACCGCCGCCGGGATCACTCTCGGCATGGACGAGCTGTACAAGGAAAGCTTTGTTAAAAAAGACCAAGCAGAACCACTACACCGAAAATTCGAACGACAATGAGGTACCGTCGACCAACTTTATTATACAAAGTTGGCATTATAAAAAAGCATTGCTTATCAATTTGTTGCAACGAACAGGTCACTATCAGTCAAAATAAAATCATTA-3’).

Subsequently the sequence was subcloned into a pDestTol2pA5 plasmid (kind gift from Chiou-Hwa Yuhm Lab) according to manufacturer’s instructions (ThermoScientific) along with p5E-Ubi (zebrafish ubiquitin promoter, kind gift from Antonio Jacinto’s Lab) and pME-mCherry (kind gift from Chiou-Hwa Yuhm Lab).

### Lentiviral plasmids for protein knock-down

All lentiviral particles were produced by co-transfection of pLenti6 and pMD 2.G (VSV-G protein) and psPAX2 (Rev and Pol proteins) into a producer cell line 293STAR RDPro (ATCC). Recombinant viral particles were harvested 48 days later, cleared for cell debris by centrifugation at 3200 g for 10 min and used. For TSG101, VPS4b, CD63, Alix, pLKO.1 plasmids with shRNA sequences were obtained from the RNAi Consortium (Broad Institute, Boston). Controls used were a non-targeting sequence (Mission; 5ʹ-CAACAAGATGAAGAGCACCAA-3ʹ; Sigma). The target sequences used were: TSG101 (KD1: 5’-GCCTTATAGAGGTAATACATA-3’; KD2: 5’-CGTCCTATTTCGGCATCCTAT-3’); VPS4b (KD1: 5-GCTGATCCTAACCATCTTGTA-3’; KD2: 5’-CCATTGTTATAGAACGACCAA-3’); CD63 (KD1: 5’-TGGGATTAATTTCAACGAGAA-3’; KD2: 5’-GCTGGCTATGTGTTTAGAGAT-3’); Alix (KD1: 5’-GCAGAACAGAACCTGGATAAT-3’; KD2: 5’-GCATCTCGCTATGATGAATAT-3’). For Syntenin-1 target pLKO.1 plasmids with shRNA sequences were obtained from Sigma mission library (KD1: 5’-CCTATCCCTCACGATGGAAAT-3’; KD2: 5’-GAGAAGATTACCATGACCATT-3’).

### Adenoviral Rab27 and RAB31 miRNA

For the knock-downs, Rab27a and Rab27b, as well as Rab31 micro-RNA (miRNA)-expressing vectors were constructed by inserting specific nucleotide sequences into pcDNA6.2-GW/EmGFP-miR plasmid harboring Pol II promoter obtained from ThermoScientific. These sequences are fused with GFP coding sequence. The synthesized oligonucleotides were annealed and ligated into pcDNA6.2-GW/EmGFP-miR according to manufacturers’ instructions. For Rab27a and Rab27b tandem miRNA sequences were transferred into pAd adenoviral vector from Thermo Scientific using Gateway technology. Rab27a miRNA1: 5’-AAACTTTGCTCATTTGTCAGG-3’; Rab27a miRNA2: 5’-TTAACTGATCCGTAGAGGCAT-3’; Rab27b miRNA1: 5’-ATTGACTTCCCTCTGATCTGG-3’; Rab27b miRNA2: 5’-TTTCCCTGAAGATCCATTCGG-3’. For Rab31 miRNA sequences were transferred into pAd adenoviral vector from Thermo Scientific using Gateway technology. miRNA1: 5’-TTTCTTTGCAGGAAACGTCCC-3’; miRNA2: 5’-TAAACTGAAGGCCATGTTGCG-3’. Viral particles were produced and cells were infected as described before ^49^.

### Transmission electron microscopy (TEM)

For ultrathin cryosectioning, cells were fixed in 2% PFA, 0.2% glutaraldehyde in 0.1M phosphate buffer pH 7.4. Cells were processed for ultracryomicrotomy and contrasted as described ^51^. All samples were examined with a FEI Tecnai Spirit electron microscope (ThermoFisher Scientific), and digital acquisitions were made with a numeric camera (Quemesa; EMSIS).

For exosomal samples, exosomes were fixed with 2% Paraformaldehyde (PFA) and deposited on Formvar-carbon coated grids (TAAB Laboratories). Samples were washed with PBS and fixed with 1% GA for 5 min. After washing with distilled water, grids were contrasted with uranyl-oxalate pH 7, for 5 min, and transferred to methyl-cellulose-uranyl acetate, for 10 min on ice. Observations were carried out using a Tecnai G2 Spirit BioTwin electron microscope (FEI) at 80 kV.

### Exosome Isolation

Exosomes derived from 40 million cultured cells were isolated from 20 mL conditioned medium. Cells were cultured in exosome-depleted medium for 48h, prepared accordingly to Lässer and colleagues ^52^. After incubation, the medium was collected and exosomes were isolated by ultracentrifugation, as described before ^52^. Briefly, the harvested supernatant was subjected to differential centrifugation at 4°C, starting with a centrifugation at 300g, for 10 min followed by a centrifugation at 16,500g, for 20 min. From this pellet, the microvesicles fraction was collected. To remove larger particles, the supernatant was filtered with a 0.22µm filter unit, after which it was ultracentrifuged at 120,000g, for 70 min, using a SW32Ti rotor. The resulting pellet was washed with PBS, and after ultracentrifugation, exosomes were resuspended in PBS.

For Mass Spectrometry (MS) an extra step of purification was introduced using a 30% sucrose cushion (Adapted from Théry. C,^53^) to eliminate sample contaminants. Exosomal pellet were resuspended in 30ml of cold PBS, loaded on top of 4 ml of Tris/sucrose/D2O solution and ultracentrifuged for 75 min at 100,000 × g, 4◦C. ∼3.5 ml of the Tris/sucrose/D2O cushion, containing the exosomes were collected using a syringe with a 18-G needle. Exosomes were then transferred to a new ultracentrifuge tube, diluted in 35 ml of PBS and centrifuged for overnight at 100,000 × g, 4◦C. The resulting pellet was resuspended in 100 µl PBS and protein quantification was performed.

### Western Blot

Cell extracts and extracellular vesicles samples were denaturated in Laemmli buffer, heated at 95°C, for 5 min. Total protein was resolved on a 10% SDS gel and transferred to a nitrocellulose membrane during 75 min at 100V. The membranes were blocked with Tris buffered saline (TBS) with 0.1% Tween 20 (TBST) containing 5% skim milk, followed by incubation with primary antibodies, overnight at 4°C. Membranes were incubated HRP-conjugated secondary antibodies for 1h at room temperature. Images were acquired using ChemiDoc Touch System (BioRad).

### Nanosight tracking analysis

Isolated exosomes were subjecte to Nanosight tracking analysis (NTA) analysis, using a NanoSight LM 10 instrument (NanoSight Ltd). Settings were optimized and kept constant between samples. Each video was analyzed to give mean, mode, median and estimated concentration for each particle size. Data were processed using NTA 2.2 analytical software.

### Optiprep linear gradient

Continuous density gradients were performed as described previously with adaptations. Briefly, 8×10^6^ cells cultured in 10cm culture dish, were washed in cold PBS and resuspended in working solution (0.25 M sucrose, 4 mM MgCl2, 8.4 mM CaCl2, 10 mM EGTA, 50 mM Hepes-NaOH at pH 7.0 with Protease Inhibitor Cocktail). Cells were lysed by sonication (3 rounds of 3 seconds at 4°C) and centrifuged at 1,000g for 5 min (2x). Continuous 5–30% optiprep gradients were prepared in working solution using a gradient mixer. Postnuclear supernatant was added to the top of 9 ml gradients and centrifuged at 100,000g for 16h using a 70.1 Ti rotor. Sequential fractions were collected (1mL each, from the top to the bottom of the tube). Fractions were recovered by ultracentrifugation at 100,000g for 30 min in a SW32Ti rotor. For Western blot, fractions were collected in PBS and further denaturated in Laemmli buffer.

### Isolation of Early and Late Endosomes by Sucrose Density Gradient Centrifugation

For the separation of early and late endosomes a protocol described by Araujo et.al was used ^26^. 25×10^6^ ARPE-19 cells, cultured in DMEM with 10% FBS, were washed with ice-cold PBS and collected with a cell scraper. Cells were collected to a 2mL microfuge tube and centrifuged 300g for 5 min at 4 °C. Cell pellet was loosen using coldfinger in a Homogeneation buffer (HB – 250mM sucrose, 3mM Imidazole, pH 7.4, 1mM EDTA, 0.03mM Cycloheximide, 10mM iodoacetamide, 2mM PMSF, 1x Cocktail inhibitor from Sigma-Aldrich). Samples were centrifuged at 1300g for 10 min at 4 °C. Supernatant was discarded, and the pellet was gently resuspended with wide-cut tip, in three times the pellet volume of HB. The suspension was passed through a 25-gauge needle, attached to a 1mL syringe, 10 times.

The homogenate was then diluted in HB (1 part homogenate to 0.7 parts HB) and centrifuged at 1,600g for 10 min at 4 °C. The supernatant was collected and centrifuged again. The postnuclear supernatant (PNS) that originate from the second centrifugation is used for organelle isolation. 5% of the PNS is centrifuged at 4°C for 10 minutes at 16,000g. The recovered supernatant was collected for cytoplasmatic fraction, free of vesicles. For the remaining PNS, the sucrose concentration was adjusted to 40.6%. The PNS was then loaded at the bottom of an ultracentrifuge tube. The 40.6% solution was overlaid with 1.5 volumes of 35% sucrose solution, 1 volume of 25% sucrose solution, and the tube was filled to the top with HB. The gradient was centrifuged at 210,000g for 16h, at 4 °C, using a 70.1Ti rotor. Each interface was collected. Late endosomes are found in the 25%/HB interface. Early endosomes are present in the 35%/25% interface. The endosomal fractions were diluted in 35 mL of HB solution and centrifuged at 100,000g for 1 h in a SW32Ti rotor. The organelle pellets were resuspended in an appropriate buffer depending on the downstream procedures.

For Western blot, fractions were collected in PBS and denaturated in Laemmli buffer. For *in vitro* uptake assays, fractions were collected in MOPS buffer and protein concentration was determined. For endosome immunoprecipitation endosomes were collected in KPBS buffer according to protocol ^54^.

### *In vitro* uptake of GST and GST-HIF1A assay

Recombinant proteins GST (Sicgen), GST-HIF1A (Abnova, H00003091-P01) were incubated with 5μg of freshly isolated vesicles and recombinant protein HSC70 (Enzo Life Sciences, ADI-SPP-751) in MOPS buffer (10 mM 3-(N-morpholino) propanesulfonic acid (MOPS), pH 7.3, 0.3 M sucrose, 5.4μM Cysteine, 1mM DTT) and ATP Regeneration Buffer (1mM ATP, 1mM MgCl_2_, 7.5mM Phosphocreatine, 50μg/mL Creatine Kinase) for 45 min at 37°C. After the incubation period, samples were cooled on ice for 1 min followed by a 30 min incubation with Trypsin at 37°C. Samples were put back to ice and denaturated in Laemmli Buffer, heated at 95°C, for 5 min and resolved on a 10% SDS-PAGE gel. The gel was transferred to a nitrocellulose membrane during 75 min at 100V. The membranes were blocked with TBS-T containing 5% skim milk, followed by incubation with antibodies against the proteins of interest.

### Rapid immunofluorescence for PAmCherry

Cells were grown on glass coverslips and fixed with 4% PFA (w/v) in PBS for 5 min at 37°C. Cells were then permeabilized with 0.05% saponin in PBS for 5 min at room temperature, washed with PBS and incubated with 50 µl primary antibody diluted in PBS containing 5% (v/v) FBS and 0.1% (w/v) BSA for 10 min at room temperature. The cells were then washed and incubated with secondary fluorochrome-conjugated antibodies diluted in PBS containing 5% (v/v) FBS and 0.1% (w/v) BSA for 10 min at room temperature. The cells were washed and viewed on the Zeiss LSM980 Airyscan confocal microscope. For Rab5QL imaging, cells were transfected with a Rab5QL fused to GFP and cells were incubated with antibody raised against one of the extracellular loops of CD63 for 30 minutes before fixation for immunofluorescence. The cells were washed and viewed on the Zeiss LSM710 confocal microscope using a x63 1.4 Plan-Apochromat oil-immersion objective. For 3D reconstruction of the images the Imaris software was used.

### Live cell imaging

Cells were culture in μ-Slide 8 well chambered coverslips (Ibidi). As previously described ^23^, lysosomes were loaded with 0.5 mg/ml of dextran 647 in DMEM with 10% FBS for 4h at 37°C followed by incubation in conjugate-free medium for 20 hr. For endosomes, the same cells were incubated with 0.5 mg/ml of dextran 488 in DMEM with 10% FBS for 10 min, followed by a chase of 10 min in conjugate-free medium. PAmCherry was photoactivated with a 405 nm laser for 1 minute. Live cells were imaged in a Zeiss LSM 710 confocal microscope using a x63 1.4 Plan-Apochromat oil-immersion objective. The images were acquired 37°C under 5% CO_2_.

### Immunoprecipitation

Cells were collected from dishes with ice cold PBS using a cell scrapper and centrifuged at 15,000g for 10 minutes. In all cases pellets (cells and exosomes) were resuspended in 150μl of lysis buffer (50mM TRIS-HCl pH 7.4, 150mM NaCl, 10mM iodoacetamide, 2mM PMSF, protease inhibitor cocktail and 0.5% of NP-40) and sonicated 3 times, 1s each at 4°C. After, samples were centrifuged at 15,000g for 10minutes and pellets were discarded. All samples were incubated with 2μg of the antibody against the protein of interest overnight at 4°C. Subsequently 30μl of protein G–Sepharose was added to the sample and incubations proceeded at 4°C for 2h. Beads were washed 3 times with lysis buffer containing 0.15% of NP-40, denatured with Laemmli buffer and boiled at 95°C for 5 min. Samples were then analyzed by SDS-PAGE. The membranes were blocked with 5% non-fat milk in TBS-T and probed for the proteins of interest.

### Endosomal immunoprecipitation

After endosomal immunoprecipitation was adapted from protocol ^54^. Briefly, endosomes were gently resuspended in KPBS (136 mM KCl, 10 mM KH2PO4, pH 7.25 was adjusted with KOH). Magnetic beads were incubated with anti-LAMP2A or LAMP2B antibodies for one hour. Subsequently the endosomes were incubated with the magnetic beads for an additional hour at 4°C. Immunoprecipitates were then gently washed three times with KPBS. The samples were denatured with Laemmli buffer and boiled at 95°C for 5 min. Samples were then analyzed by SDS-PAGE. The membranes were blocked with 5% non-fat milk in TBS-T and probed for the proteins of interest.

### Subcellular fractioning

To evaluate the presence of exosomal HIF1A in the nuclei of receiving cells, exosomes were isolated from ARPE-19 cells that were under hypoxia with 300μM of CoCl_2_, for 12h. Subsequently 5×10^5^ 769-P cells were incubated with 5μg of exosomes for 1h. Cells were collected and fractioned. Briefly, cells were collected with a cell scraper, with ice-cold PBS. Cells were centrifuged for 10 min at 300g. Cell pellet was resuspended in 50μl of lysis buffer for 30 min, on ice (50mM TRIS-HCl pH 7.4, 150mM NaCl, 10mM iodoacetamide, 2mM PMSF, 1x Cocktail inhibitor from Sigma-Aldrich). Samples were then centrifuged at 4°C for 10 minutes at 1,000g. Supernatants were removed to new 1.5 mL microfuge tube and pellet (nuclear pellet) was lysed with Laemmli buffer. Supernatant was centrifuged at 4°C for 10 minutes at 16,000g. The resultant pellet (vesicular pellet) and the second supernatant (cytoplasm) were also lysed with Laemmli buffer. Samples were then analyzed by SDS-PAGE. The membranes were blocked with 5% non-fat milk in TBS-T and probed for the proteins of interest.

### Reporter assay

HIF1A and HSF1 activity was measured by transducing cells with the reporter gene Luciferase under the control of either the Hypoxia Response Element (HRE) or the Heat Shock Element (HSE), respectively, using ONE-Glo™ Luciferase Assay System (Promega), according to the manufacturer’s specifications.

### Zebrafish *in vivo* experiments

A zebrafish lines were maintained in a circulating system with 14 hour/day and 10 hour/night cycle periods at 28°C. The mating and spawning of zebrafish were incited by the change of dark to light. Embryos were collected before they start independent feeding, at 5 days postfertilization (dpf). Therefore no ethical approval was necessary according to the Council Directive 2010/63/EU on the protection of animals used for scientific purposes ^55^. Embryos were maintained at 28°C in E3 medium. For live imaging, *casper*, *Tg(kdrl:mCherry) or Tg(mpeg1:mCherry)* embryos, at 100 cell stage, were injected in the YSL with pUbi-mCherry-p2A-GFP-ExoSignal in the presence or absence of Morpholionos at a 150 nm concentration, with a microinjector under a stereomicroscope. We used a Syntenin-a morpholino oligonucleotide directed against the translation start site (5’-ACAACGACATCCTTTCTGCTTTCA-3’^35^), LAMP2A morpholino oligonucleotide directed against the LAMP2A unique intron-exon junction (5’-AGCTGAAAATAAAGAGAATGAGTGA-3’) and a Standard Control morpholino oligonucleotide (5’-CCTCCTACCTCAGTTACAATTTATA-3’). Morpholinos were obtained from Gene Tools LLC (Philomath, OR). At 3 dpf, embryos were anaesthetized in buffered 200 μg/mL MS-222 (Sigma) and incubated 1 h with 300 nm Bafilomycin A before being immobilized in a 1% low melting and imaged in by Spinning Disk or Airyscan Confocal (Zeiss LSM980). For neovascularization, assays at 3 dpf 0,25μg of exosomes was injected into the Duct of Cuvier of *Tg(fli1a:EGFP)* zebrafish with a microinjector under a stereomicroscope. At 5dpf, embryos were fixed with 4%PFA, overnight, at 4°C. Embryos were washed in PBS and mounted in Glycerol Mounting Media. Images were acquired in a Zeiss LSM 710 confocal microscope using a x40 1.2 C-Apochromat Water-immersion objective.

When required optical slices were acquired and 3D reconstruction of the images and quantifications were performed using Imaris software.

For exosome isolation 20-40 zebrafish larvae were dissociated at 28°C using Liberase for 30 min with gentle agitation. Subsequently exosomes were isolated from the samples as previously described.

### Statistical analysis

Data are reported as the means ± standard deviation of at least three independent experiments. Comparisons between multiple groups were performed by one-way analysis of variance test (ANOVA) with Tukey’s multiple comparison tests, using GraphPad Prism 8.0 software (GraphPad Software). For comparison between two groups, the paired t-test was used. In all cases, p < 0.05 was considered significant.

## Supporting information

Graphical Abstract

## Figure Legends

**Supplemental Figure 1. Isolated sEVs are enriched in exosomal markers while LAMP2A KO does not interfere with endosomal morphology.** A) ARPE-19 cells were transduced using lentiviral particles containing vectors for the expression of spCas9 protein and for the gRNA directed to the CT terminal of the LAMP2A isoform. Cells were grown in colonies until reaching around 50 cells per colony. Colonies were screened for the presence of LAMP2A by Western Blot. Western blot of whole cell lysates with antibodies against LAMP2, LAMP2A, LAMP2B and Actin (ACTB). LAMP2A KO does not affect the protein levels of the LAMP2B isoform. B) Cells were prepared according to the indicated protocols. Electron micrographs of WT and LAMP2A KO ARPE-19 cells show endosomes containing ILVs with the same morphology. C) ARPE-19 cells were cultured in exosome-depleted medium. sEVs were isolated from cell culture supernatants. MS based analysis of sEVs show that the isolated vesicles are enriched in sEVs markers.

**Supplemental Figure 2. mCherry-ExoSignal presence in sEVs.** ARPE-19 cells were cultured in exosome-depleted medium. sEVs were isolated from cell culture supernatants. A) Isolated sEVs were separated in a discontinuous sucrose gradient. Recovered fractions were blotted with antibodies raised against CD63 and mCherry. Results show that mCherry-Exosignal is present in vesicles from sucrose fractions that are enriched in the markers CD63 with densities that are typical of exosomes. B) Isolated sEVs were incubated with trypsin in the presence or absence of 1% of Triton X100. mCherry, unlike the membrane protein Cx43, is resistant to trypsin, indicating that mCherry-Exosignal is located in the inside of the sEVs. C) Quantification of Rab27 depletion.

**Supplemental Figure 3. The PAmCherry-ExoSignal chimeric protein is sorted into early endocytic vesicles.** WT and KO LAMP2A ARPE-19 cells were transduced with lentiviral particles containing vectors for the expression of PAmCherry-ExoSignal. A) PAmCherry-ExoSignal was photoactivated with 405 nm laser for 1 min. Live cells were imaged in a Zeiss LSM 710 confocal microscope. Lysosomes are labelled in yellow, endosomes in cyan and PAmCherry-ExoSignal in magenta. In WT cells, arrows at the 5 min time point show that endosomal compartments are negative for PAmCherry-ExoSignal. At the 15 min time point, on the other hand, endocytic vesicles are positive for PAmCherry-ExoSignal, and both 647 and 488 dextrans, indicating that early endosomal compartments are already fusing with lysosomes. In LAMP2A KO cells PAmCherry-ExoSignal puncta in residual. B) Cells were transfected with Rab5QL-GFP and incubated for 30 min with antibody against CD63 extracellular loop for 30min before fixation. Immunofluorescence using confocal microscopy shows that PAmCherry-ExoSignal puncta localizes in Rab5QL-GFP compartments, inside CD63 positive ILVs when LAMP2A is present. 3D images were reconstructed using Imaris software.

**Supplemental Figure 4. Hypoxia Inducible Factor 1A (HIF1A) is present in sEVs and endosomal fractions.** A) WT and LAMP2A KO ARPE-19 cells were cultured in exosome-depleted medium and incubated with 300 μM of the hypoxia-mimetic agent CoCl2 for 12h. Western Blot of cell extracts and exosomal fractions using antibodies raised against LAMP2A, HIF1A, FLOT1 and CD63 show that HIF1A is absent from LAMP2A KO exosomes. 769-P cells were transduced with lentiviral particles containing WTHIF1A and KFERQ-mutatant HIF1A. Cells were cultured in exosome-depleted medium for 12h. Western Blot of cell extracts and exosomal fractions using antibodies raised against TSG101, Alix and HIF1A show that mutated HIF1A is not present in exosomes. C) WT and KO LAMP2A cells was loaded on top of a continuous 5–30% optiprep gradient. Nine sequential fractions were collected. Western blot of isolate fractions with antibodies raised against endosomal markers CD63, FLOT1, LAMP2, LAMP2A, Alix, TSG101, Rab5 and Rab7 show HIF1A is present in endosomal fractions (7 and 8), only when LAMP2A is present. All samples were analyzed under the same experimental conditions. The results represent the mean ±SD of N=3 independent experiments (n.s. nonsignificant; *p < 0.05; **p < 0.01; ***p < 0.001).

**Supplemental Figure 5. mCherry-ExoSignal colocalizes with early endosomal markers and e-MI substrates GAPDH and Aldolase are loaded into endosomes independently of LAMP2A.** A) Immunofluorescence using LSM 980 Aryscan confocal microscopy of cells fixed in methanol with antibodies against mCherry, LAMP2A, EEA1 and m6p or Rab5-GFP. PAmCherry-ExoSignal shows high co-localization coefficient with LAMP2A, EEA1 and RAB5-GFP only when LAMP2A is present. B) GAPDH-GST and Aldolase-GST was incubated with freshly isolated vesicles in the presence/absence of ATP, the molecular chaperone HSC70, and LAMP2A. Samples were treated with trypsin to degrade all the protein that was not protected by the endosomal membrane. Western blot using antibodies raised against the GST-tag show that both proteins are translocated into both EEs and LEs enriched fractions, independently of LAMP2A.

**Supplemental Figure 6. mCherry-ExoSignal co-localizes with ESCRT-independent endosomal machinery.** A) Confocal images of methanol fixated cells show recovery of colocalization between mCherry-ExoSignal and CD63, Alix or Rab31, but not with TSG101, VPS4b and Syn-1, when LAMP2A expression is rescued in LAMP2A KO cells. B) Quantification of KD efficiency.

**Supplemental Figure 7. LAMP2A mediates loading of KFERQ-containing proteins into sEVs in the zebrafish larvae.** A) Schematic representation of zebrafish experiment using mCherry-P2A-GFP-ExoSignal. B) *casper* zebrafish at 1000 cell stage were injected at the YSL with empty vector (sham) or GFP alone. Confocal microscopy using a spinning disk shows no GFP in the zebrafish larvae caudal plexus at 3dpf. C) WB of sEVs isolated from *casper* zebrafish after injection with GFP-ExoSignal-P2A-mCherry. Only GFP-ExoSignal is present in isolated sEVs. D) sEVs were isolated from Zebrafish expressing human CD63-pHluorin injected at 1 cell stage in the presence or absence of Syntenin-a morpholino. Syntenin-a morpholino decreases sEVs levels. E) 3D reconstruction of *Tg(kdrl:mCherry)* larvae, expressing mCherry in endothelial cells, injected with sham or both GFP-ExoSignal and Syntenin-a morpholino. No GFP signal is visible in the caudal plexus. F) *Tg(mpeg1:mCherry)* larvae, expressing mCherry in macrophages, sham injected. No GFP signal is present in macrophages located in the caudal plexus. G) WT and KO LAMP2A ARPE-19 cells were cultured in exosome-depleted medium incubated with 300 μM of the hypoxia-mimetic agent CoCl_2_ for 12h before exosome isolation from media supernatants. 769-P cells were incubated with the WT derived exosomes for 1h. Subsequently, cells were harvested and fractioned into Nuclear fraction (N), Cytoplasm fraction (C) and Vesicular fraction (V). Western Blot of the fractions using antibodies raised against HIF1A, Tubulin (TUB) and Lamin B, show that exosomal HIF1A is reaches the nucleus of the receiving cell. All samples were analyzed under the same experimental conditions. The results represent the mean ±SD of at least N=3 independent experiments (n.s. nonsignificant; *p < 0.05; **p < 0.01; ***p < 0.001).

**Supplemental Figure 8. Schematic representation of the proposed mechanism for cytosolic protein loading into exosomes.** Cytosolic proteins containing KFERQ-like motifs are recognized by the chaperone HSC70 and targeted to the endosomal membrane, where it binds to the receptor LAMP2A. As the endosomal membrane invaginates to create ILVs, proteins are trapped in their lumen while binding to LAMP2A. The LAMP2A-mediated loading of proteins is assisted by additional molecular machinery, including CD63, Alix, Syntenin-1, Rab31 and ceramides, rather than ESCRT molecular components such as TSG101 and VPS4b. Subsequently, endosomes filled with ILVs travel to the cell periphery, in a process likely mediated by Rab27, fusing with the plasma membrane and releasing its contents as exosomes.

## Acknowledgements

This work is supported by Portuguese Foundation for Science and Technology (JVF grant: SFRH/BPD/121271/2016, ARS grant: PD/BD/106052/2015, CM grant: PD/BD/136902/2018, PP grant: PTDC/SAU-ORG/118694/2010) and iNOVA4Health Research Unit (LISBOA-01-0145-FEDER-007344), which is cofunded by Fundação para a Ciência e Tecnologia / Ministério da Ciência e do Ensino Superior, through national funds, and by FEDER under the PT2020 Partnership Agreement; https://www.fct.pt/index.phtml.en, http://www.inova4health.com. The funders had no role in study design, data collection and analysis, decision to publish, or preparation of the manuscript. We would like to thank CEDOC’s Fish and Microscopy Facilities, as well as, IMM Fish Facility for their invaluable help. Additionally, we would like to acknowledge Chiou-Hwa Yuhm’s Lab and Antonio Jacinto’s Lab for their help in developing the zebrafish tools, as well as, Bruno Costa Silva’s Lab for the help with NanoSight Tracking analysis.

