## Supplementary figures and images for "LAMP2A regulates the loading of proteins into exosomes"

### Graphical Abstract

#### Graphical Abstract

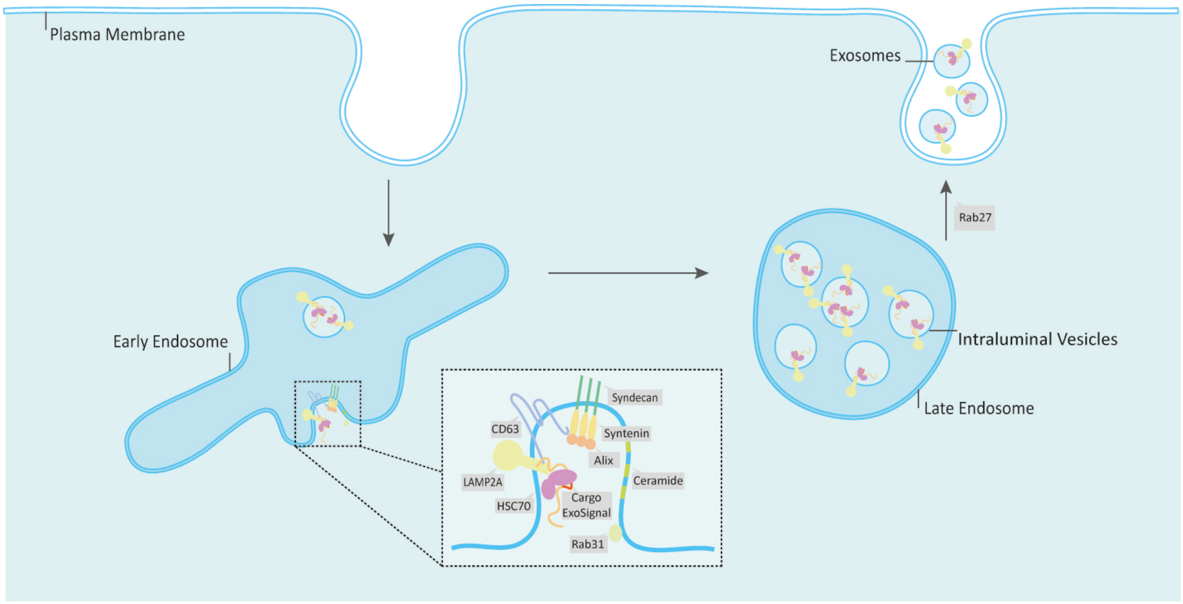

### Supplemental Figure 1

#### Supplemental Figure 1

A

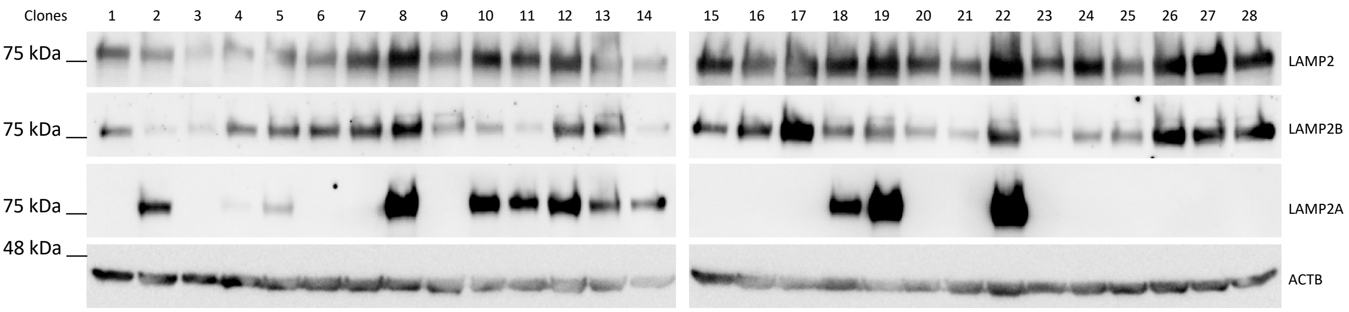


B

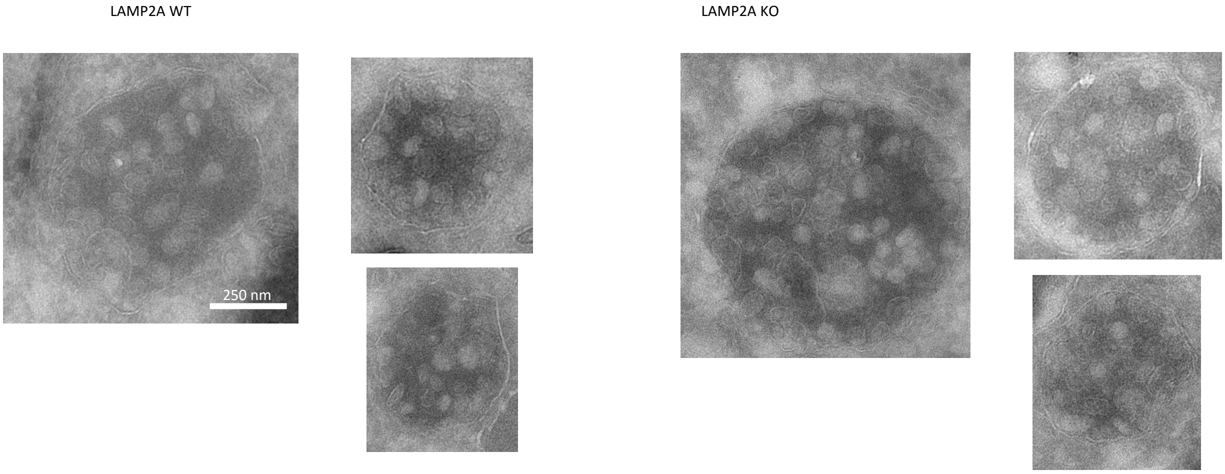


C

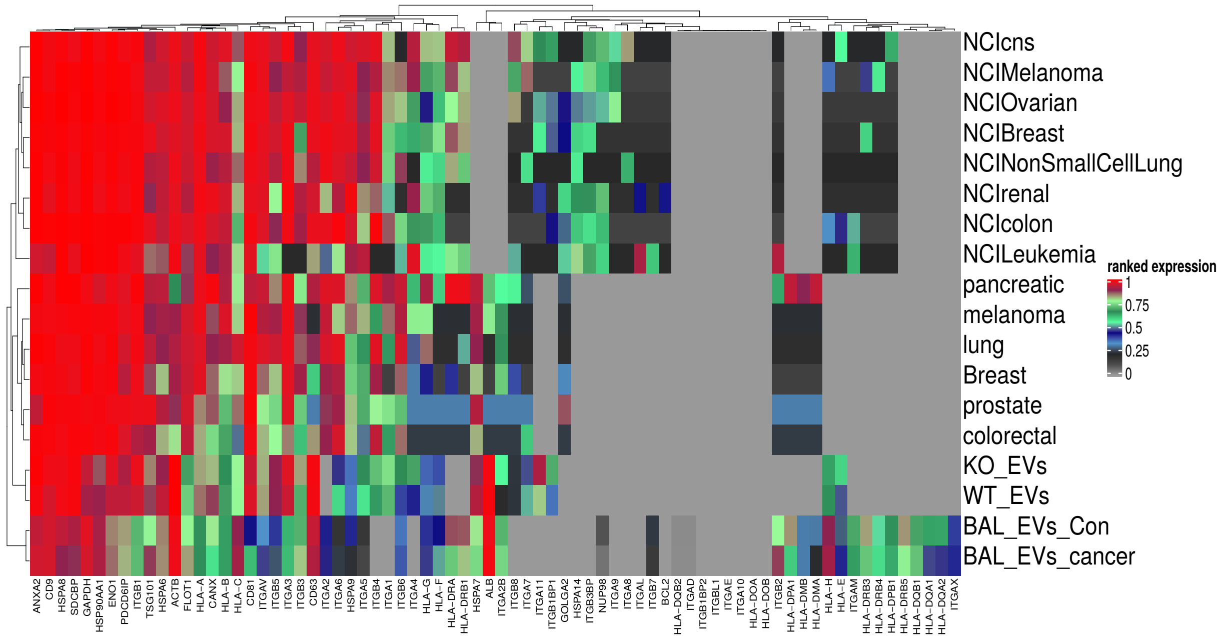

### Supplemental Figure 2

Supplemental Figure 2

A

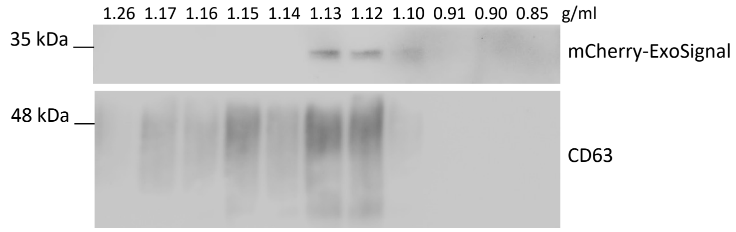


B

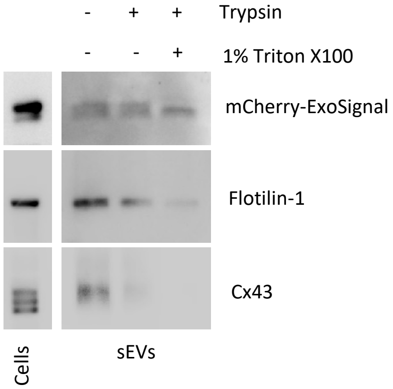


C

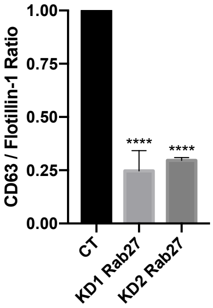

### Supplemental Figure 3

Supplemental Figure 3

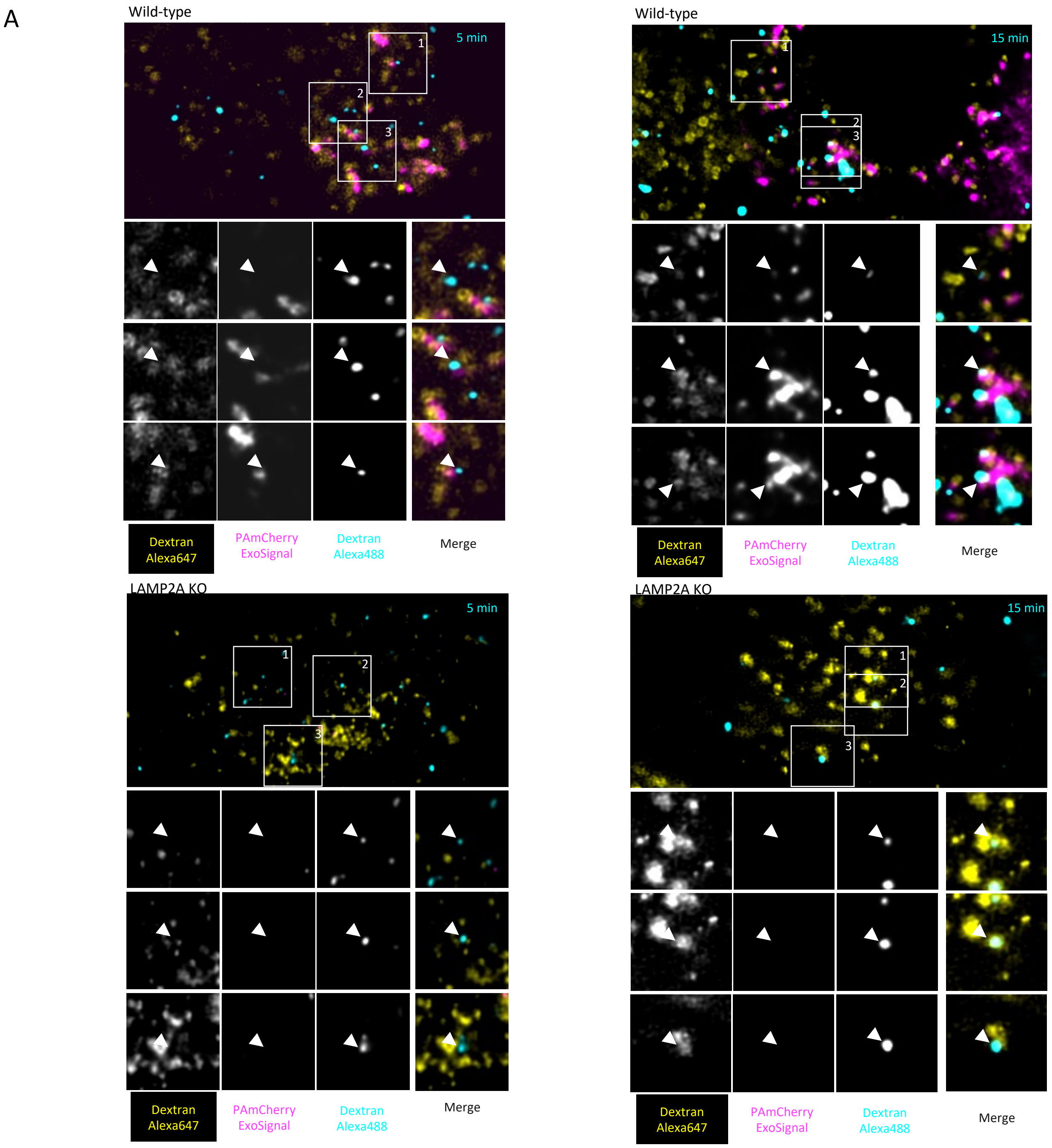


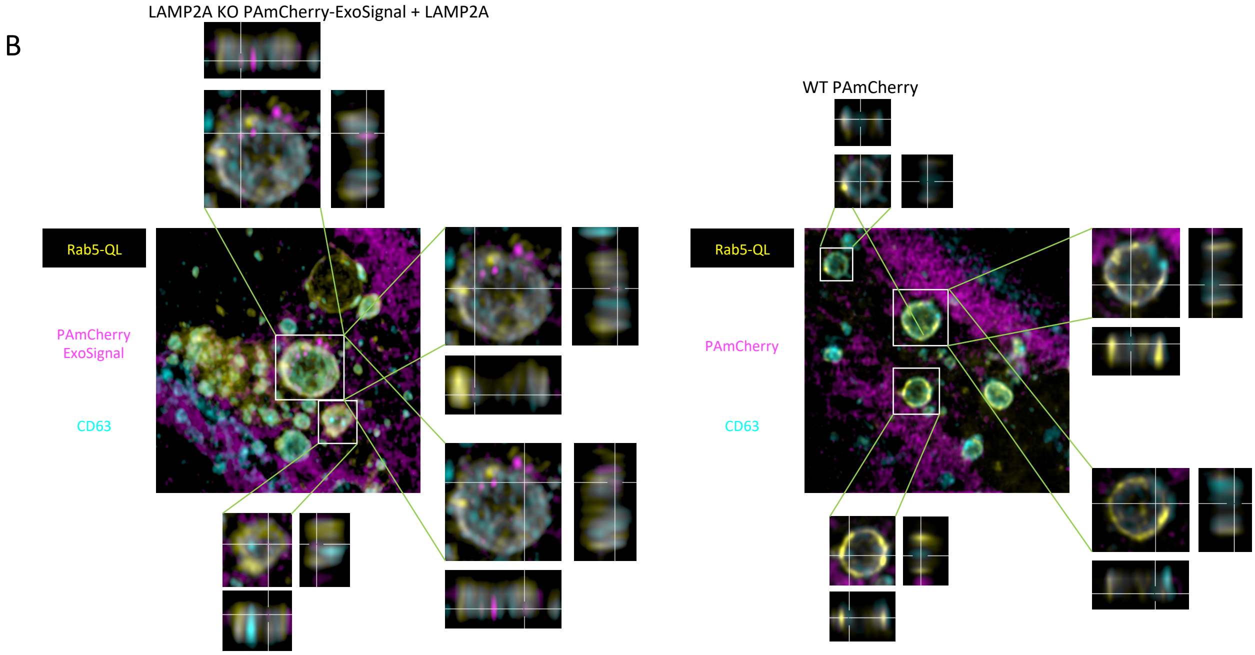

### Supplemental Figure 4

Supplemental Figure 4

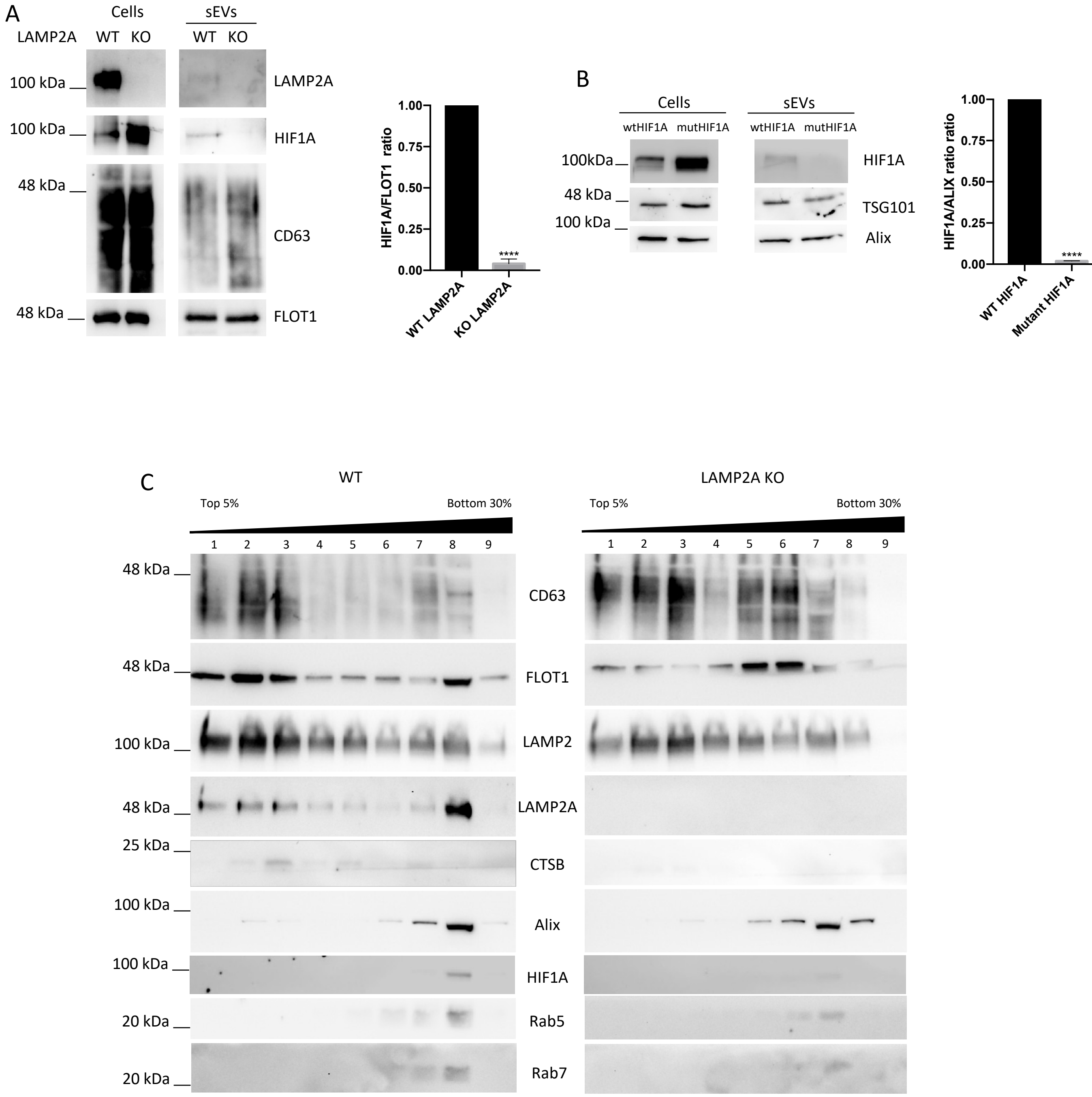

### Supplemental Figure 5

Supplemental Figure 5

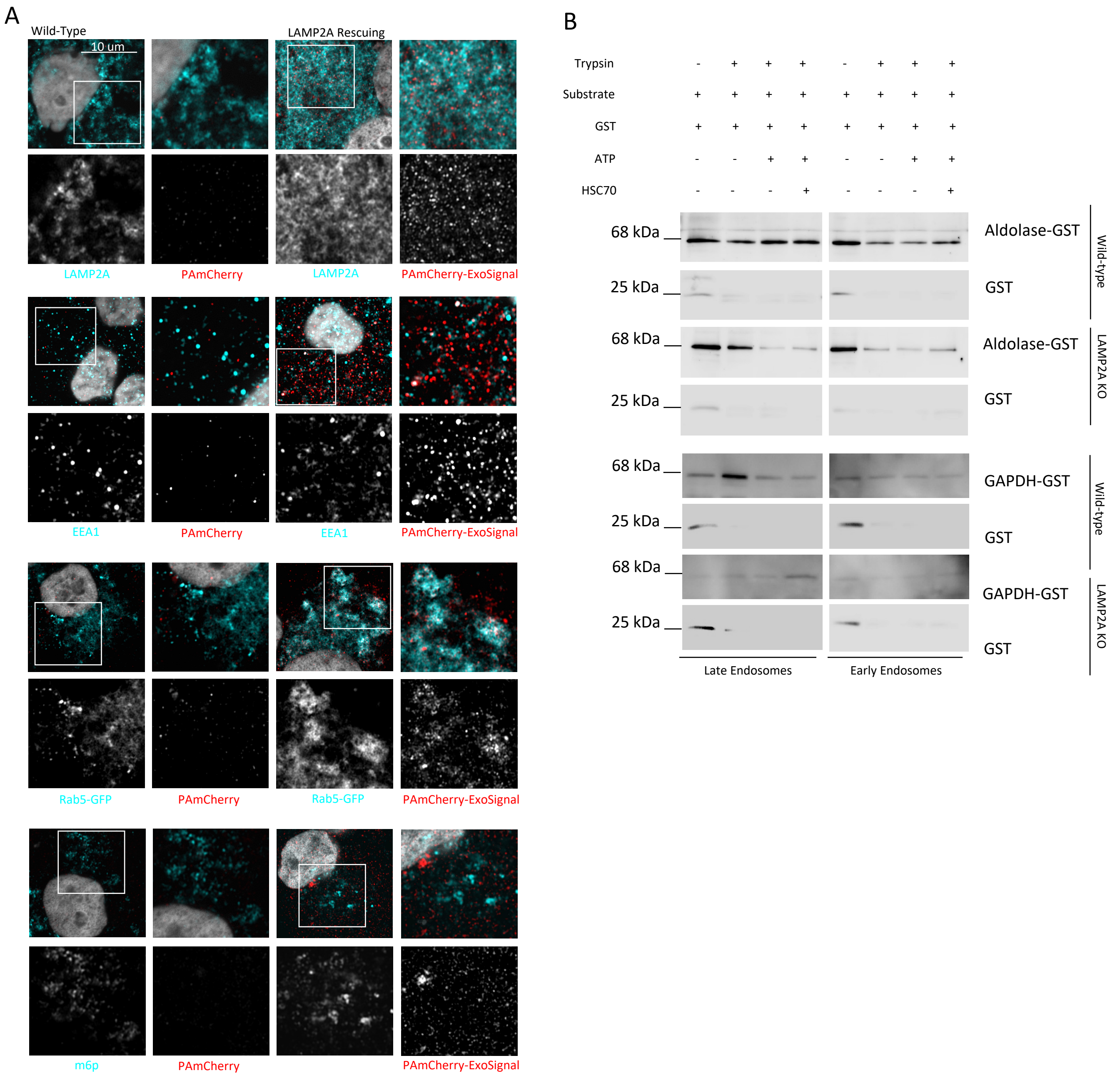


m6p

### Supplemental Figure 6

Supplemental Figure 6

A

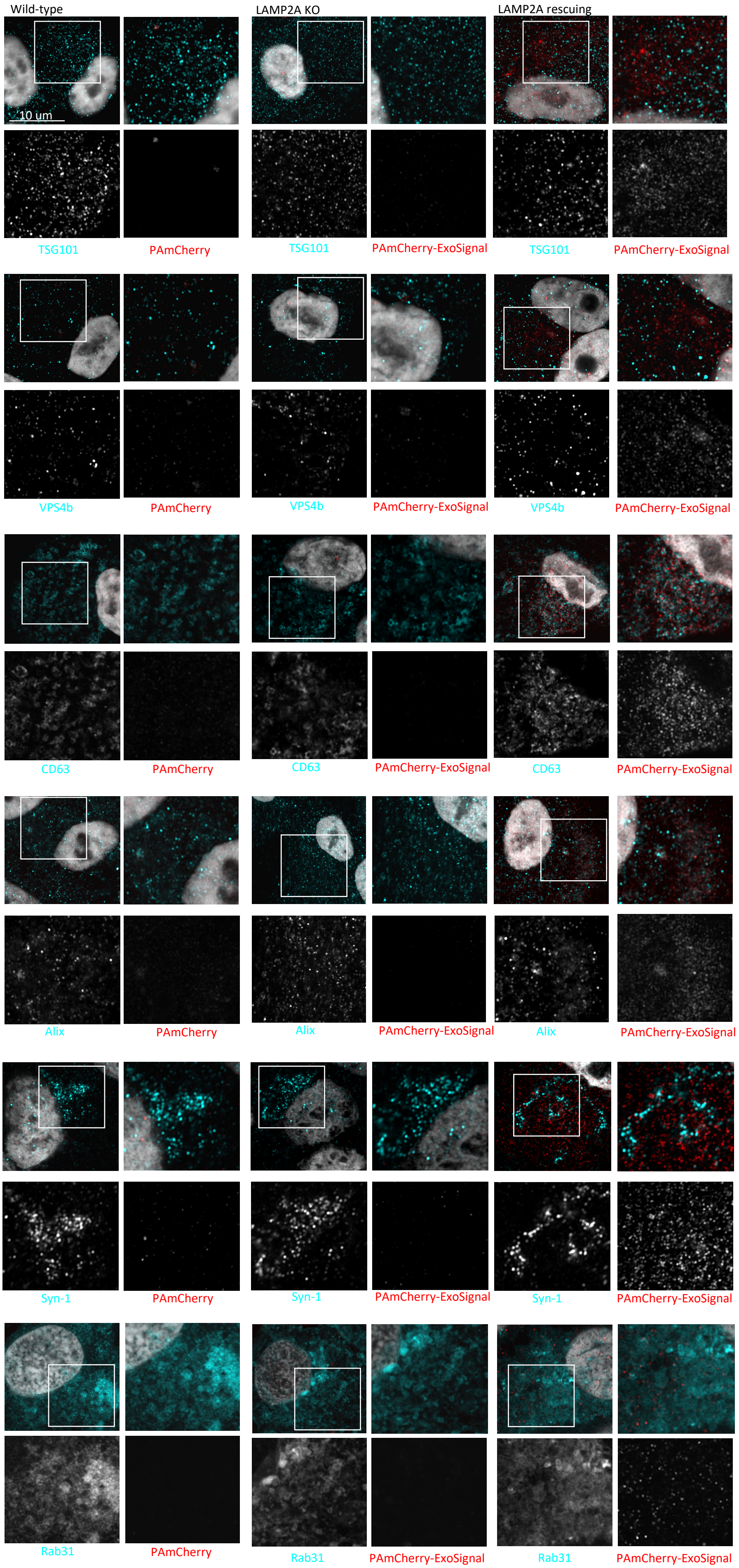


B

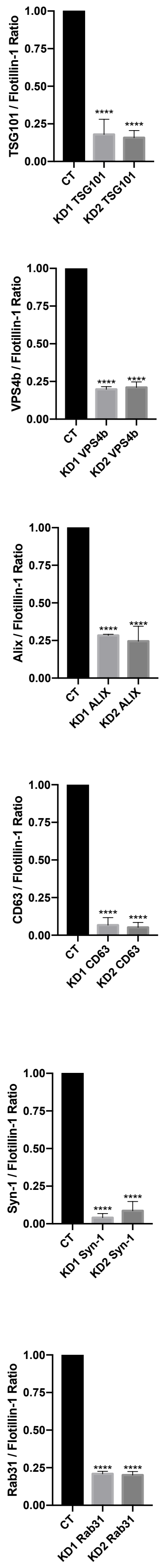

### Supplemental Figure 7

#### Supplemental Figure 7

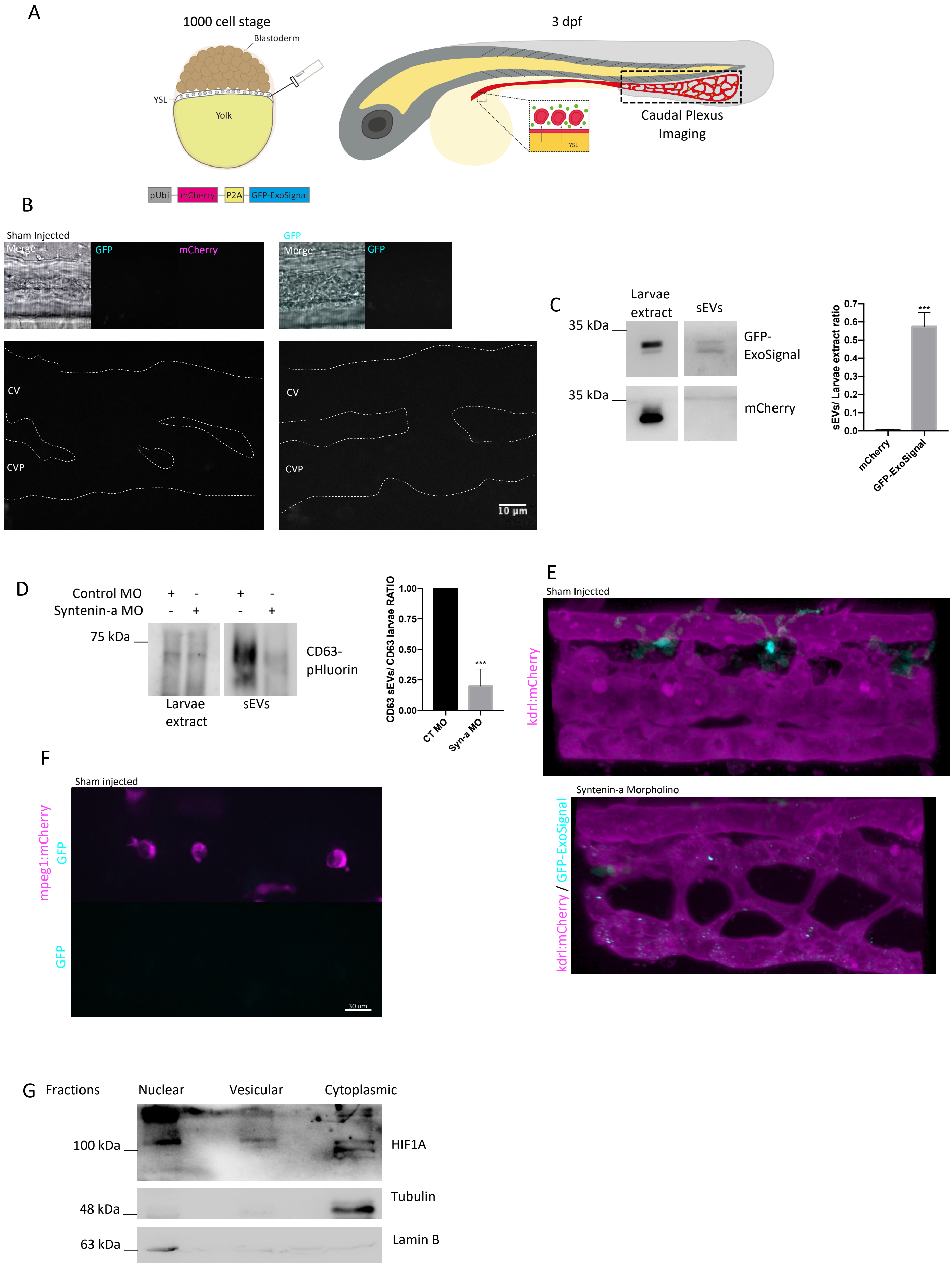
